# Establishment of a novel fetal ovine heart cell line by spontaneous cell fusion

**DOI:** 10.1101/2022.07.14.500071

**Authors:** Khalid M. Suleiman, Mutaib M. Aljulidan, Gamal eldin M. Hussein, Habib N. Alkhalaf

## Abstract

We established a unique immortal cell line designated, FOH-SA, by serial passage of fetal ovine heart cells. In a novel phenomenon in cell biology, we demonstrated that the immortalization of the line supervened as a result of spontaneous cellular and nuclear fusion of two morphologically distinct cardiocytes in passage 29. Fused cells gave progeny cells which grew into multicellular filaments. Trypsinization of the filamentous multicellular growth gave the immortal heart cell line.

Comparative single nucleotide polymorphism (SNP) genotyping of the cell line before and after cell fusion revealed a large scale genetic conversion resulting in 96% homozygosity in SNPs genotypes in the progeny cells. Partial sequencing of the mitochondrial (mt) genome of the cell line revealed the occurrence of large mutational events in the control region, the *tRNA-Phe* and *12S rRNA* genes of the mt genome of progeny cells. The cell line was found permissive to sheep pox, peste des petits ruminants (PPR), lumpy skin disease (LSD), rift valley fever (RVF), and camel pox viruses.

This study would resolve the over half a century mystery of how the African Green Monkey cell line (VERO) had evolved. We authenticated the cell line at the European Collection of Authenticated Cell Cultures (ECACC) and deposited it at the American Type Culture Collection (ATCC) for the purpose of patenting under the Budapest Treaty.

## Introduction

Cell culture became indispensible tool in the field of modern biotechnology and has been used in production of human and veterinary viral vaccine, therapeutic recombinant proteins, interferons, and monoclonal antibodies [1, 2, 3]. It has also been used to study intracellular reactions, as an *in vitro* model for research, the study of cytotoxicity of pharmaceuticals and bacterial toxins [4, 5] and recently cell culture emerged as future candidate for cell-based meat for human consumption [6].

Primary cells obtained from mature or embryonic human and animal organs usually undergo a limited number of passages after which they enter replicative senescence [7, 8], characterized by degenerative changes such cell rounding, enlargement, decreased capacity to proliferate, and detachment from the monolayer. At the molecular level senescent cells show the distinctive markers of senescence such as increased expression of p53, p21, p16 and other cyclin-dependent kinase inhibitors such as p27 and p15 [9,10,11].

In rare cases some mammalian cells under continuous subculturing escape replicative senescence and become spontaneously immortal resulting in a continuous cell line [12, 13]. The Madin-Darby bovine kidney (MDBK) and the Madin-Darby ovine kidney (MDOK) cell lines were spontaneously immortalized by serial passage of cells from renal tissue of *Bos Taurus* and *Ovis aries* respectively [14]. Recently, an ovine kidney cell line (FLK-N3) was also spontaneously established by serial passage of primary fetal lamb kidney cells from a fetus of a normal sheep [15]. The ubiquitous African green monkey cell line (VERO) was also spontaneously immortalized by serial passage of kidney cells of it was deposited at ATCC at passage 113 to establish a bank for availability [16].

Over the last 35 years, numerous types of primary cell cultures had been used at the Veterinary Vaccine Production Centre (VVPC), Riyadh, Saudi Arabia, to produce sheep pox vaccine, which included ovine cells from the testicles, aortic endothelium, kidneys, and heart. Although the titers of the produced vaccine were satisfactory, the disadvantages associated with primary cultures such as risk of contamination, tediousness, inconsistency of cell characteristics, the ethics of slaughtering animal and use of organs to prepare cell cultures, and the limited production capacity hurdled the production wheel especially in the face of increased demand in Saudi Arabia for sheep pox vaccine that necessitated trials over the years to establish a continuous cell line to overcome the difficulties and disadvantages associated with the use of primary cultures in vaccine production.

In this study we succeeded to establish a continuous cell line by serial subculturing of fetal ovine heart cells. In a novel finding in cell biology, *and* in a novel finding in cell biology, we demonstrated that the cell line evolved from spontaneous fusion of two distinct heart cell phenotypes.

The cell line was found highly permissive to many animal viruses and therefore it would be of importance in isolation of animal viruses, vaccine development, cell biology studies, and other biotechnological fields.

## Materials and methods

### Ethics declarations

A single pregnant ewe was used in the study and it was handled in compliance with the Implementing Regulations of the Law of Ethics of Research on Living Creatures of the National Committee of Bioethics (NCBE) for the use of animals in experiments (Article, 38). The authors confirm that all the methods used in the study were approved by the Ministry of Environment, Water and Agriculture (MEWA). The authors also confirm that all the procedures and the experimental protocols were in accordance with the regulations and guidelines of MEWA.

### Preparation of primary cell culture

In March 2013 a single pregnant ewe, *Harri* breed, was euthanized by exsanguination in accordance with the Implementing Regulations of the Law of Ethics of Research on Living Creatures of the National Committee of Bioethics (NCBE) (Article, 38) for collection of fetal organs to prepare primary cell cultures necessary for production of sheeppox vaccine. The fetal heart was aseptically incised and used to prepare primary cell cultures. The heart muscle was chopped to 0.5-1.0 cm, washed 3 times with 1x MEM (Gibco) and digested under stirring with 0.25% trypsin ten times ½ an hour each. After each cycle of digestion, the suspension was allowed to settle and the supernatant was sieved through sterile gauze and was centrifuged at 1000 RPM for 5 minutes. The supernatant was then discarded and the cell pellet was cultured into non-vented Roux flasks containing complete growth medium (CGM) consisted of 1X MEM (Gibco), 10 % fetal bovine serum (FBS) (Gibco), Glutamax 1% (Gibco), Gentamycin 50*µ*g/ml, and pH 7.2-7.4 adjusted with 5.6% sodium bicarbonate. Flasks were incubated at 37° C in a normal incubator.

### Established cell lines

Primary heart cell cultures obtained from extractions one to eight were initially subcultured after five days and then after every three days and this cell line was used for production of sheep pox vaccine. A second cell line was established from heart extractions nine and ten which was first subcultured after nine days and was then serially passaged every three days. Samples of the second heart cell line at different passages were cryopreserved in Recovery Cell Culture Freezing Medium (Gibco) in liquid nitrogen and at -80° C.

### Cell morphology and cell line transformation

Initially, the morphology of the two cell lines was studied under an inverted bright field microscope, and later, the cell morphology of the second cell line (extractions nine and ten) was studied under an inverted phase contrast microscope.

Cryopreserved cells passages 20 to 27 were resuscitated and subcultured to study the transformation event under a phase contrast microscope (Olympus CKX53) with a camera (DP74, software CellSens). Cultured flasks were tape fixed on the microscope stage in a walk-in normal incubator at 37°C. We photographed fixed fields of passages 28, 29, 30, 32, and 33 every three hours for 24 hours and then every 24 hour for a total of 72 hours for each passage.

### Growth Curve and population doubling time

A three-day old cultures of heart cell line passages 25 (before transformation, BT) and 36 (after transformation, AT) were seeded into 25 cm^2^ non-vented Roux flasks at a density of 9144 cell/ cm^2^ and a density of 6485 cell/ cm^2^ respectively. Flasks were incubated at 37° C in a normal incubator and cells were counted after day 1, 2, 4, 6, 8, and 10 using Handheld Automated Cell Counter (Millipore). The growth curve of the two passages was constructed by plotting the log of growth against time and the population doubling time was calculated using standard formula [17].

### Storage of the cell line at 37° C

We observed that the monolayer of the cell line established after transformation (passage 33 and above) could be maintained at 37° C for 3-4 weeks without showing signs of cell degeneration. To confirm this observation, we incubated cultures of passage 59 at 37° C for 6 months with the growth medium being changed every three weeks. During storage, the monolayer was examined under the inverted microscope every week. At the end of the storage period flasks were subcultured and the sensitivity of the cells to sheep pox virus was determined.

### DNA preparation

DNA was extracted from a cell suspension containing approximately 5x 10^5^ cell/ ml using MagNA Pure 96 DNA and Viral NA Small Volume Kit in Magna pure 96 as instructed by the manufacture (Roche, Germany). DNA concentration was measured by NanoDrop 2000 instrument (Thermo Scientific, 260 nm). DNA extracted from cells in passages 22, 47, and 59 was aliquoted and stored at -80° C.

### Authentication and mycoplasma testing

A DNA sample of passage 22 was shipped to European Collection of Authenticated Cell Cultures (ECACC), UK, for authentication of the cell line by mitochondrial DNA barcoding and detection of mycoplasma contamination. Currently, DNA barcoding is used to authenticate animal cell lines and it utilizes the mitochondrial cytochrome c oxidase (COXI) subunit I gene as a barcode. The test is performed by PCR amplification of a 617 bp segment of the mitochondrial COXI gene followed by sequencing the product and matching it to a library of reference sequences [18].

### Single nucleotide polymorphism (SNP) genotyping

We shipped Aliquots of DNA samples of cell at passage 22 (BT) and 47 (AT) to NEOGEN®’s GeneSeek laboratory, USA, for SNP genotyping using the Ovine SNP50 BeadChip. The OvineSNP50 Beadchip a high throughput microarray system developed by Illumina in collaboration with the International Sheep Genomics Consortium (ISGS). The bead chip utilizes the Infinium® HD Assay to analyze the DNA samples. Briefly, the infinium assay work-flow consisted of an initial PCR-free amplification of the DNA samples in day one, followed by enzymatic fragmentation of the amplified samples, then application of fragmented samples to the bead chip, and incubation of the chip overnight for samples to hybridize on day two. On day three, hybridized samples were extended, fluorescently stained and imaged by the iScan® System.

### Partial sequencing of the cell line mitochondrial genome

We sequenced two segments of the FOH-SA cell line mitochondrial genome of passages 26 (BT) and 59 (AT) using Sanger Sequencing with 3730xl DNA Analyzer and BigDye Terminator v3.1 Cycle Sequencing Kit (Applied Biosystems, USA). Two pairs of primers designed by Meadows [19] from the complete ovine mtDNA [20] were used to amplify the segments. The first pair of primers CytB-F 5’ GTCATCATCATTCTCACATGGAATC-3’ and CytB-R5’ CTCCTTCTCTGGTTTACAAGACCAG -3’ was for amplification a 1272 bp region of the ovine mitochondrial cytochrome b gene (AF010406 positions 14078 to 15349). The second pair of primers, mtCR-F2 5’ AACTGCTTGACCGTACATAGTA-3’ and mtCR-R1 5’-AGAAGGGTATAAAGCACCGCC-3’ was used to amplify a 1246 bp fragment spanning part of the control region, the complete sequence of *tRNA-Phe* gene, and partial sequence of *12S rRNA* coding RNA gene (AF010406 positions 15983 to 592).

### Susceptibility of the cell line to viruses

The sensitivity of the FOH-SA cell line to viruses was investigated by infecting it with several animal viral vaccines’ stains which included, sheep pox virus (Romanian strain), Peste des petits ruminants (PPR) (Nigerian 75/1), Rift valley fever (RVF) (Smithburn strain), Lumpy skin disease (LSD) (Neethling strain), and a local attenuated isolate of camel pox virus. Passage 36 cells were inoculated at multiplicity of infection (MOI) of 0.01 and flasks were incubated at 37° C and the monolayers were daily examined under an inverted microscope for development of cytopathic effect (CPE). Flasks were harvested when the CPE reached 90-95% and the tissue culture infective dose 50 (TCID50) for each virus was determined by Karber method [21].

### Production sheep pox vaccine

We have used the FOH-SA cell line since 2013 for production of sheep pox (Romania strain) and camelpox vaccines (local attenuated isolate). Cells were propagated in Roux (125, 175, 225 cm^2^) and roller flasks (850, 1700 cm^2^) and inoculation was done with multiplicity of infection (MOI) of 0.01. The viral suspensions were harvested when about 90% CPE was reached.

### Cell line deposition at ATCC for patenting

Thirty cryotubes each containing more than 10^6^ cells of the fetal heart cell line (FOH-SA) passage 51 were shipped on dry ice to the Foreign Animal Disease Diagnostic laboratory (FADDL), United States Department of Agriculture (USDA) in Plum Island for safety test to food and mouth disease (FMD), PPR, sheep pox, and camel pox viruses as a prerequisite by USDA prior to deposition at ATCC. After passing safety test, 25 cryotubes were shipped from FADDL to ATCC for deposition for the purpose of patenting under the Budapest Treaty on the International Recognition of the Deposit of Microorganisms for the Purposes of Patent Procedure.

### Data analysis

Illumina GenomeStudio software (GSGT Version 2.0.2) and GenCall Version 7.0.0 were used to call and report the genotype of SNP loci on the bead chip from sample images of DNA samples of heart cell line passage 22 and 47 generated by iScan® System from the bead chip [22].

Data of the partial sequencing of the mitochondrial genome were controlled for quality and trimmed by DNA Base Assembly (v5.15.0.) and Pairwise Sequence Alignments (PSA) of mitochondrial DNA sequences were carried out with EMBOSS Needle [23].

## Results

### Cell lines and morphology

The first cell line obtained from heart extractions one to eight consisted of fibroblast-like cells. The monolayers became confluent in five days and cells were then subcultured every three days and this cell line was used for production of sheep pox vaccine till it reached replication senescence at passage 11.

The second cell line derived heart extractions nine and ten was named Fetal Ovine Heart cell line-Saudi Arabia (FOH-SA). Initially this cell line showed scanty growth, however, it was confluent after nine days. The cell line revealed two cell morphologies, epithelial-like and fibroblast-like cells and this morphology prevailed to passage 28, then the cells became transformed in passage 29 through passage 32. Cells after passage 33 exhibited a constant epithelial morphology and were smaller in size compared to the mother cells. FOH-SA cell line was successfully subcultured every three days till passage 140.

### Transformation of the ovine heart cells

#### Bright field microscopy

During serial passage of heart cell line, a constant cell transformation event was documented under bright field microscopy which began at passage 27 by increased cell density of fibroblast-like cells. In passage 29 the two cell morphologies disappeared and replaced by filaments monolayer. Wes documented that cells in passage 29 became connected with tubules detectable three hours post incubation that led to cell fusion, the nuclei of connected cells move to a central location in the connecting tubule and became coupled to each other. Then huge amounts of mitochondria from fused cells migrated and accumulated around and the nuclei of fused cells. This was followed by separation of fused cells into two progeny cells. After 72 hrs incubation passage 29 the monolayer appeared in form of filaments. The filamentous cell morphology was also documented at passages 30, 31, and 32. At passage 33 the cell line stopped growing in filaments and thereafter, the morphology remained constant till the highest passage reached (passage 140).

The cell line that evolved after transformation consisted of one phenotype and had an increased proliferative potential and showed a different CPE pattern to sheep pox virus compared to the cells before transformation.

### Phase contrast microscopy

More details of the heart cells’ morphologies and the cell fusion process demonstrated by light microscopy were revealed by investigating the cell line under phase contrast microscopy. The fibroblast-like cells consistently produced long threads that were detectable 3 hrs post incubation and remained demonstrable after 24 hrs but became disintegrated so that it is not demonstrable after 48 hrs incubation (Fig.1). In passage 28 the two cell phenotypes appeared as islets of epithelial-like cells surrounded by the fibroblast-like cells (Fig. 2).

**Fig 1.**
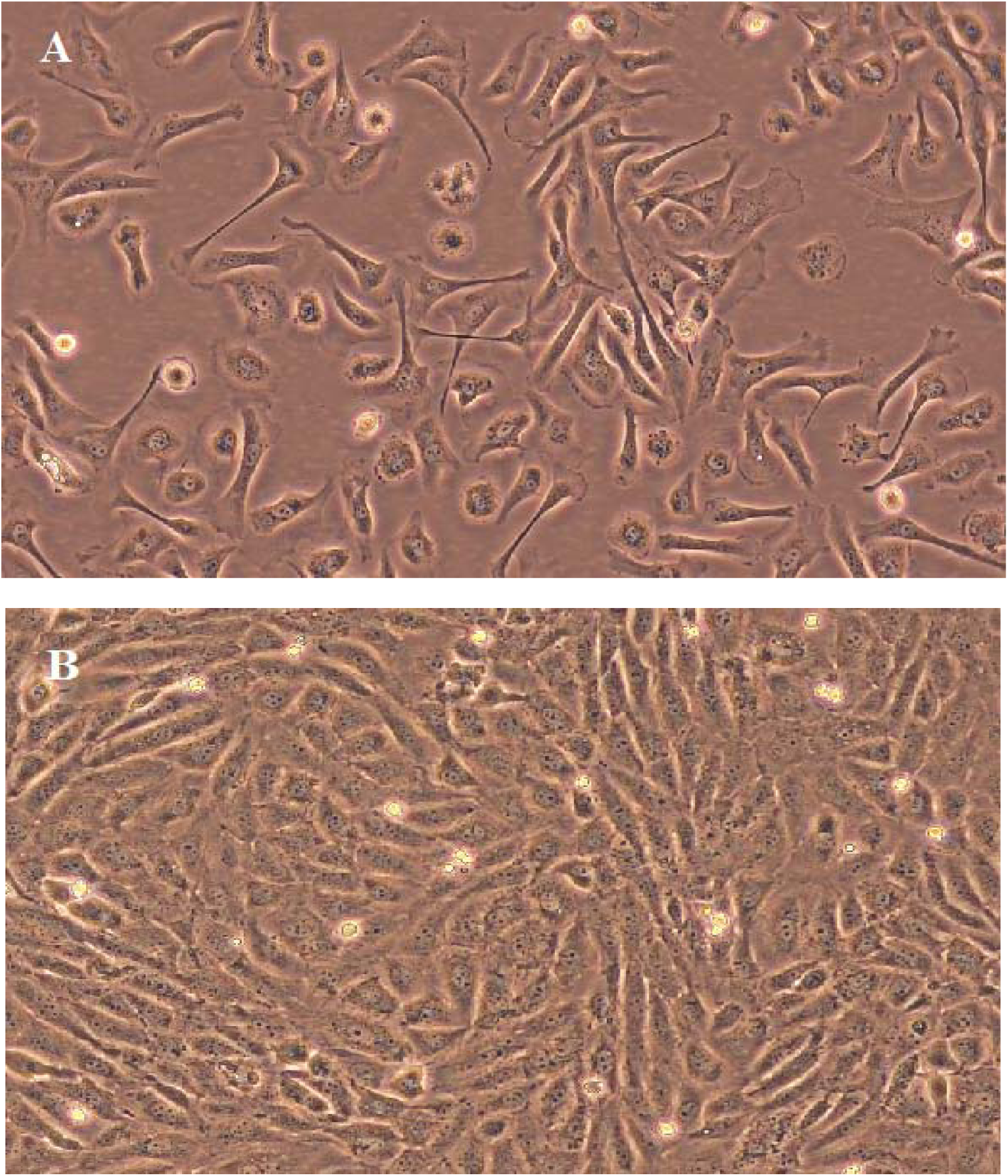
Morphology of the heart cell line passage 22. a) a three hour culture showing the fibroblast-like cells developing long threads and the epithelial like cells appearing rounded in shape, and b) the same culture in (a) after 48 hrs incubation showing both cell types with the fibroblast-like cell without threads, phase contrast X200.

**Fig 2.**
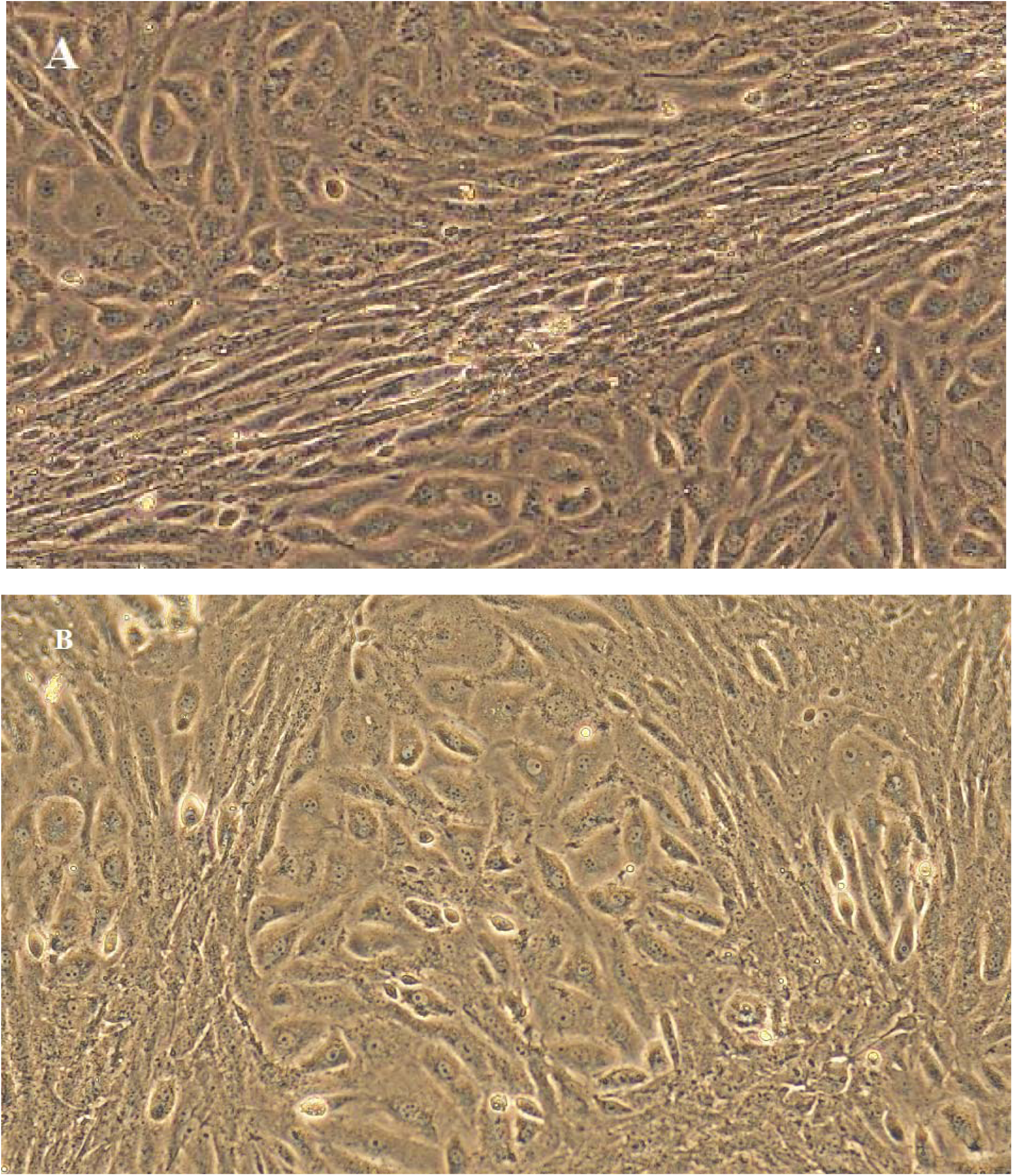
Morphology of the heart cell line. a) passage 26 showing the fibroblast-like cells in a bundle growing diagonal to field and the epithelial-like cells above and below the bundle and b) passage 28 showing the epithelial-like cells surrounded by bundles of fibroblast-like cells, phase contrast X200.

In passage 29 the two heart cell phenotypes instead of growing and multiplying, the fibroblast-like cells projected their threads reaching the epithelial-like cells resulting in establishment of connections that developed into a tubules connecting the two cell types. When the two cell phenotypes existed neighboring each other they directly fuse and such fusion points could be detected 3 hrs post incubation of passage 29 cells and the nuclei of fused cells appeared coupled together with huge amount of mitochondria accumulating around them (Fig. 3 a-c).

**Fig 3.**
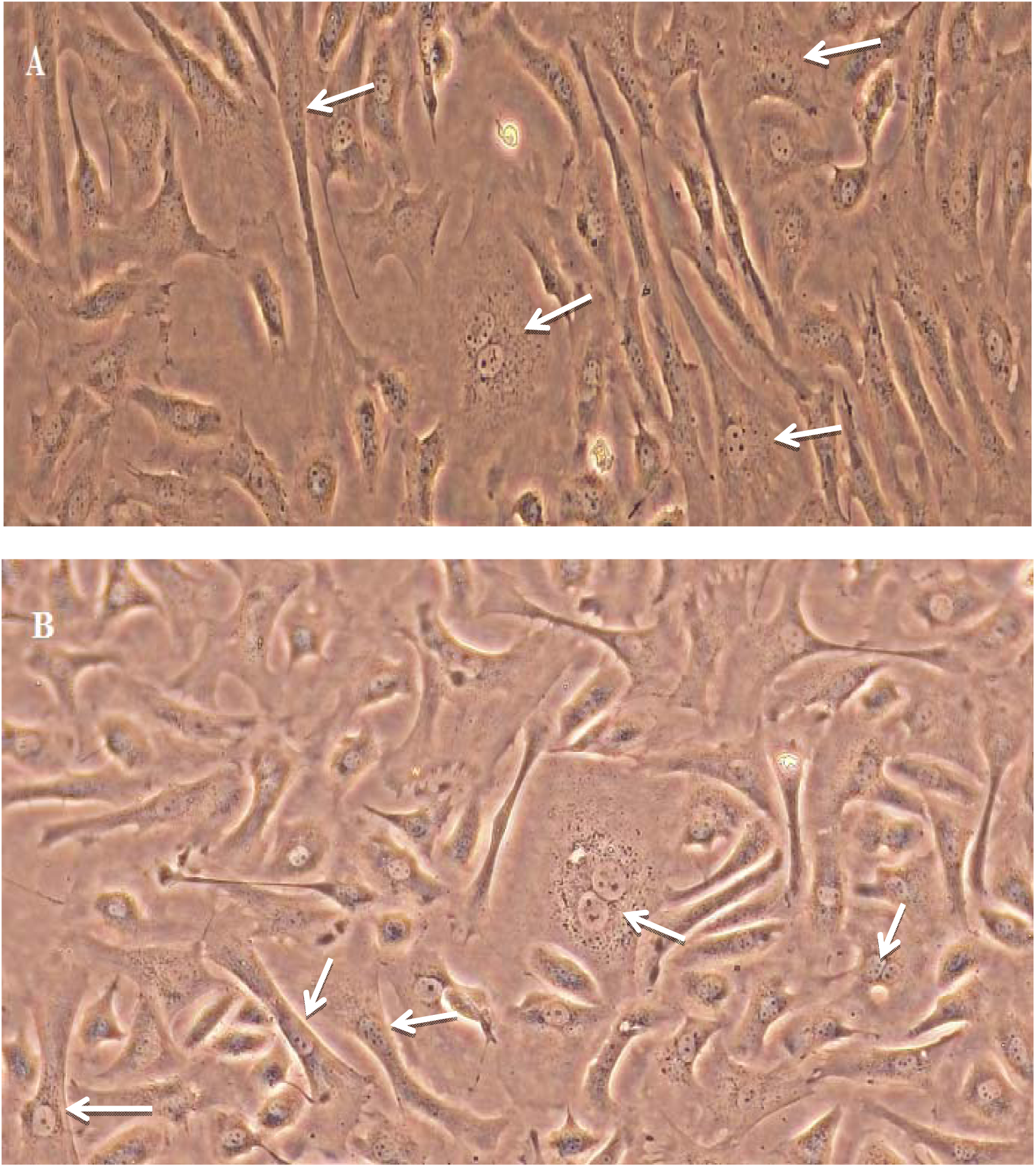
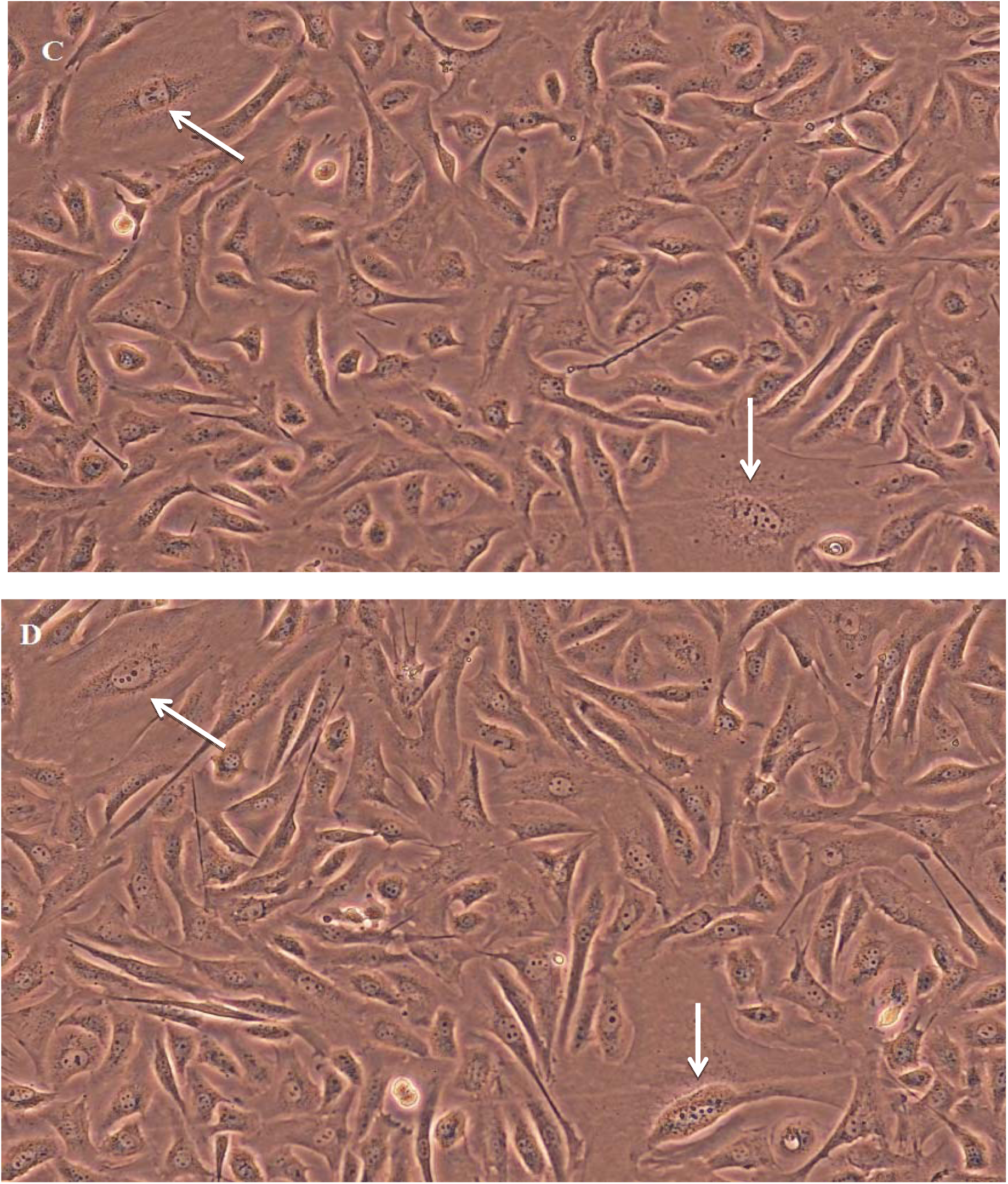
A three hour culture of the heart cell line passage 29. a), and b) Showing heart cell phenotypes connected by tubules and points of spontaneous cell fusion with aggregated mitochondria around coupled nuclei of fused cells, c) and d) showing two point of cell fusion after 3 and 6 hours incubation, phase contrast X200.

When mitochondrial recruitment was complete, the two nuclei became confined and compressed in part of the cytosol of the two fused cells and become completely delineated from the rest of the cytosol of fused cells. This was followed by a reaction in which complete fusion of the two nuclei was documented, characterized by disappearance of the nucleoli and increased brightening of the fused nuclei; we described this event as “nuclear fusion reaction”. When the nuclear fusion reaction was complete, the fused nuclei amalgamated with part of the cytosol of the fused cells to create the body of evolving progeny cells that eventually separate into two daughter cells (Fig. 4 a-f).

**Fig 4.**
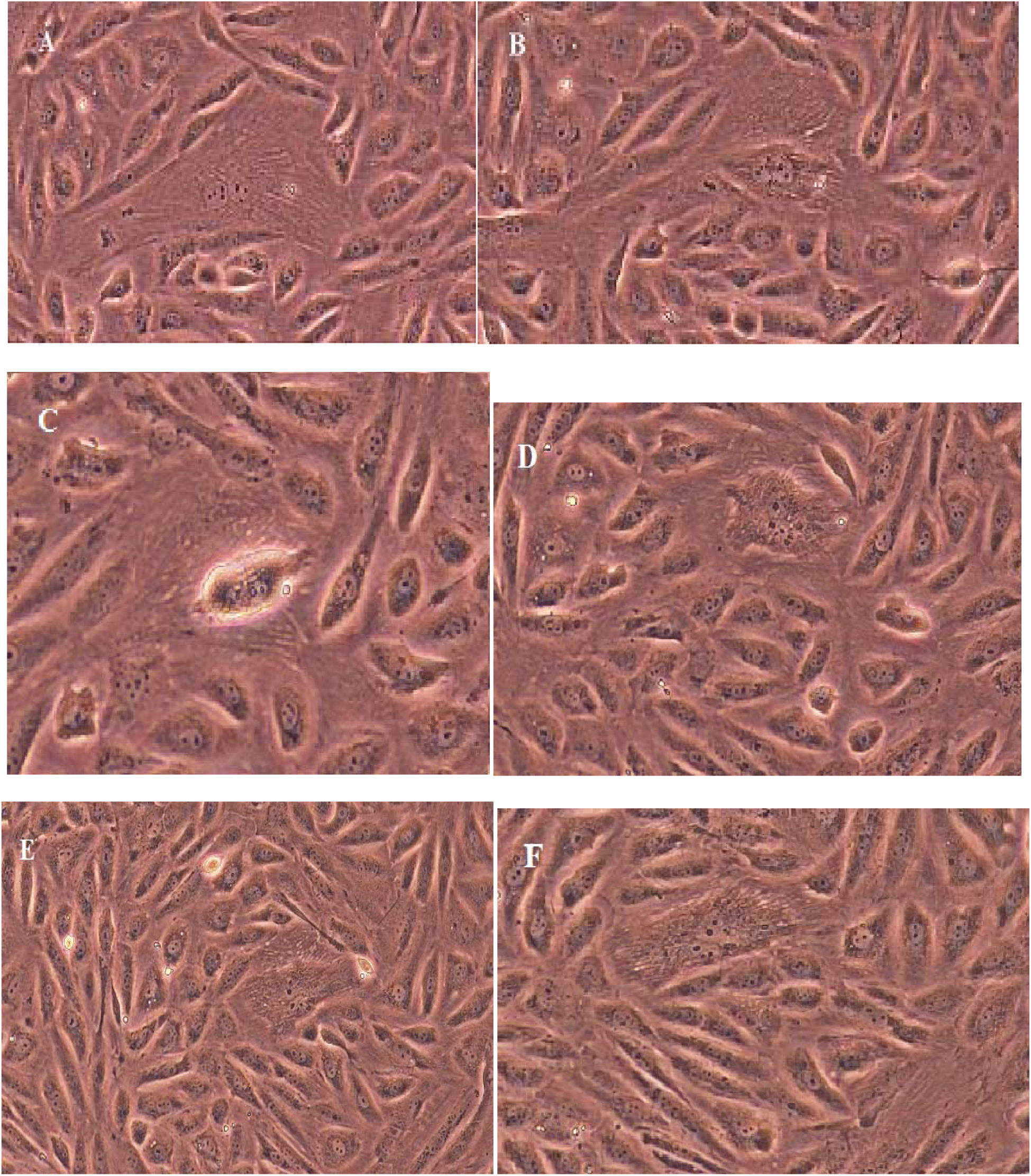
Stepwise progression of the spontaneous cell fusion process in passage 29 heart cells. a) three hours post incubation showing cell fusion and coupling of nuclei, b) 7 hours post incubation, the coupled nuclei became demarcated with part of the cytosol, c) 8 hrs post incubation, the nuclear fusion reaction, d) 11 hrs post incubation, the nucleoli of the two progeny began to appear, e) 18 hrs post incubation, the nuclei of the progeny cells became prominent, f) 21 hrs post incubation, the two nuclei of the progeny cells settled against each other, phase contrast X200.

The first progeny cells from spontaneous cell fusion came to existence in about nine hours after incubation. It was observed that horizontally coupled nuclei gave rise to nascent cells with epithelial-cell like morphology, while fibroblast-like progeny cells evolved from the vertically coupled nuclei (Fig. 5).

**Fig 5.**
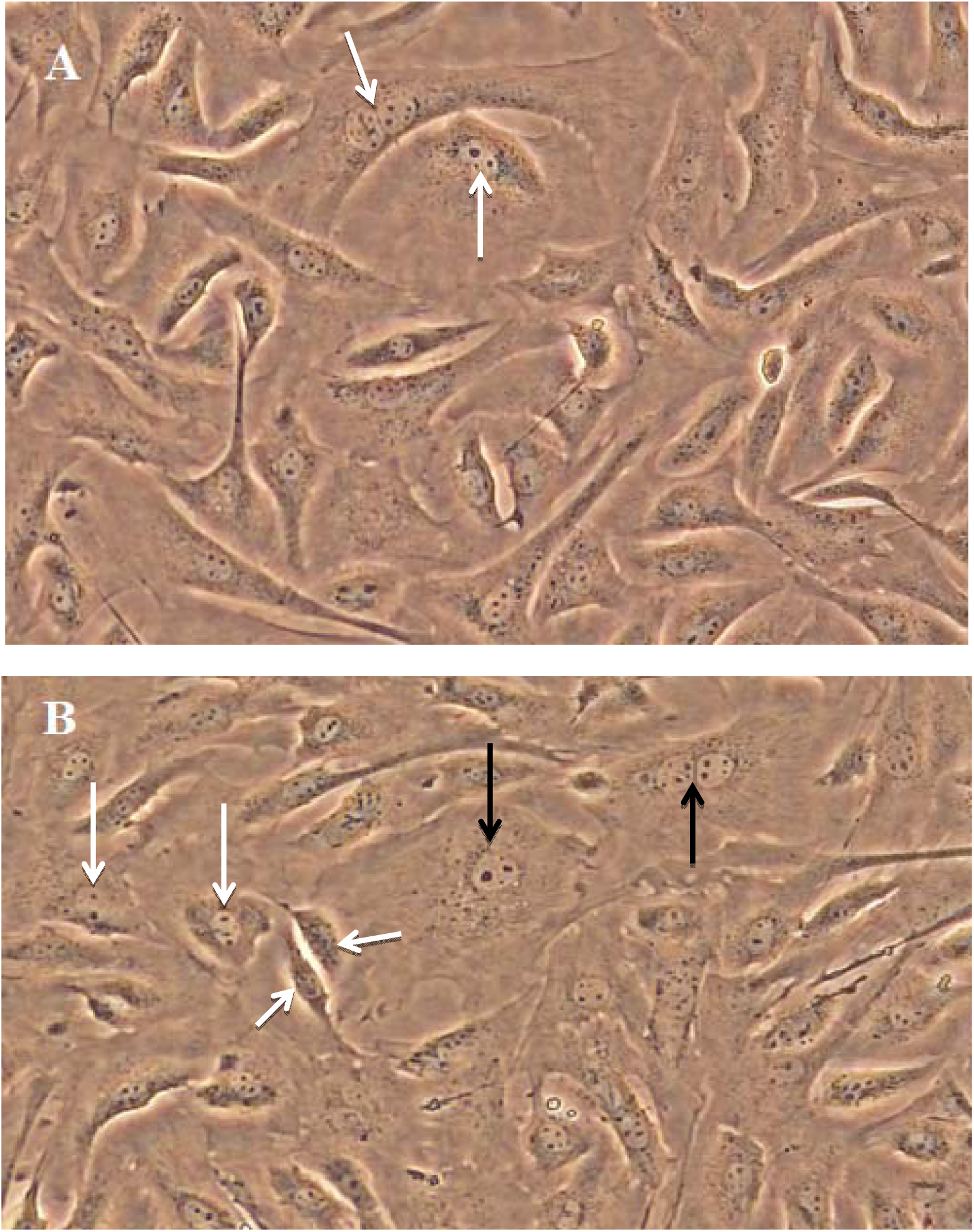
Heart cell line passage 29.a) a field six hrs post incubation showing twopoints cell fusion (white arrows) and b)The same field after 9 hrs incubation showing two fibroblast-like cells and two epithelial-like cells that developed from the two fusion points in (a) (white arrows) and in the same field two new cell fusion points came in the field (black arrows).

The process of cell fusion and the appearance of new progeny cells continued in passage 29 cells and 24 hours after incubation approximately 95% of the cell population had completed the spontaneous cell fusion and produced progeny cells (Fig. 6 a). Each progeny cell resulting from cell fusion in passage 29 then grew and multiplied so that after 72 hrs post incubation progeny cells appeared as long multi-cellular filaments (Fig. 6 b-d). Trypsinization of the 72 hour culture of passage 29 resulted in single cells (Fig. 7) that when cultured (passage 30); the cells again grew into multi-cellular filaments. The pattern of multi-cellular filamentous growth was also documented in passages 31 and 32. Finally, when the 72 hours growth passage 32 was trypsinized and cultured (passage 33), the progeny cells stopped growing into multicellular filaments and thereafter the cell morphology remained constant till the highest passage reached (passage 140) (Fig. 8).

**Fig 6.**
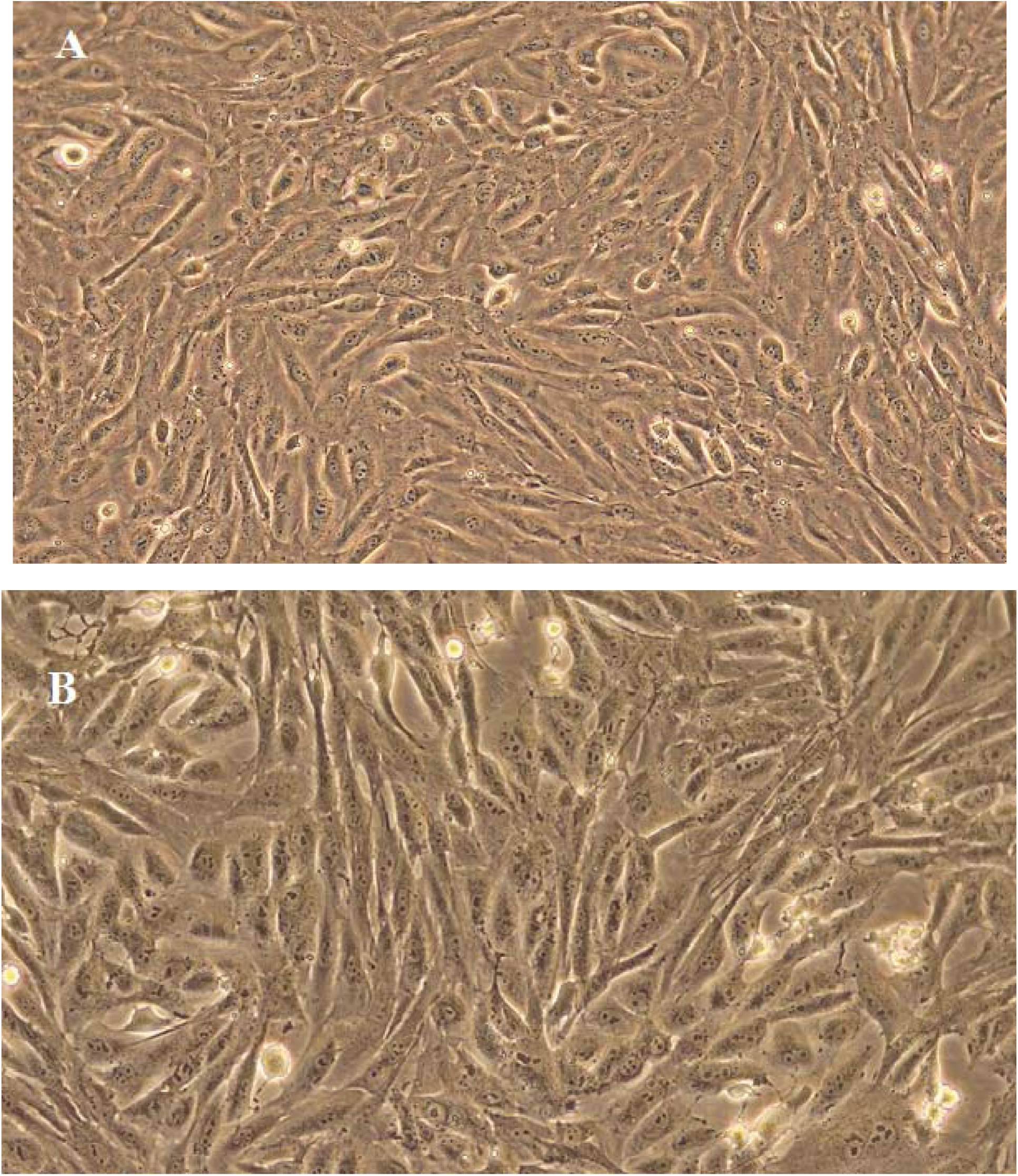
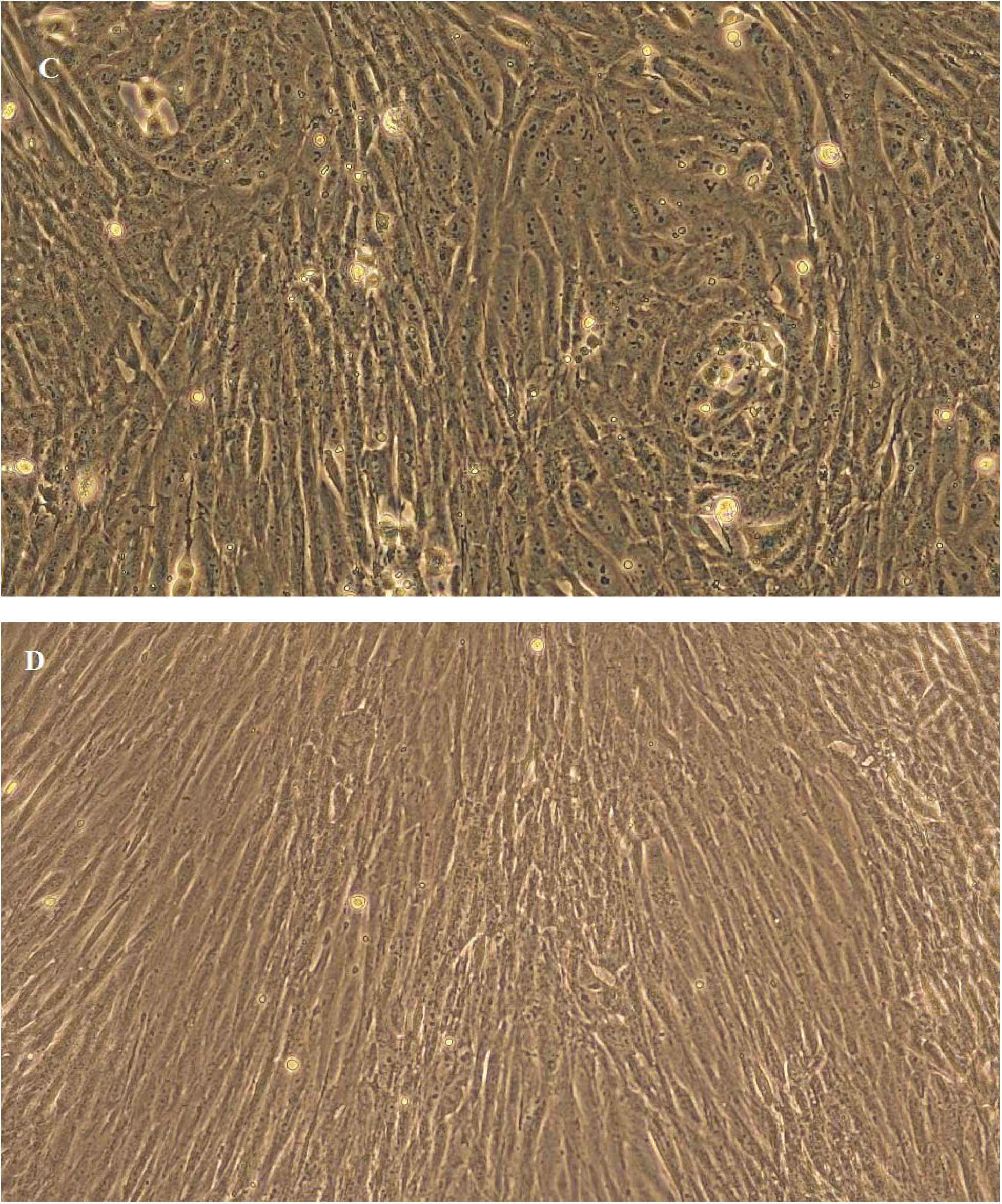
Growth of the progeny cells developed by spontaneous cell fusion in passage 29 (the same cultureof fig. 5). a) progeny cells 24 hrs post incubation, b) growing progeny cells after 36 hrs post incubation, c) 48 hrs post incubation, and d) 72 hrs post incubation the progeny cells have grown into long multicellular filaments, phase contrast X200.

**Fig 7.**
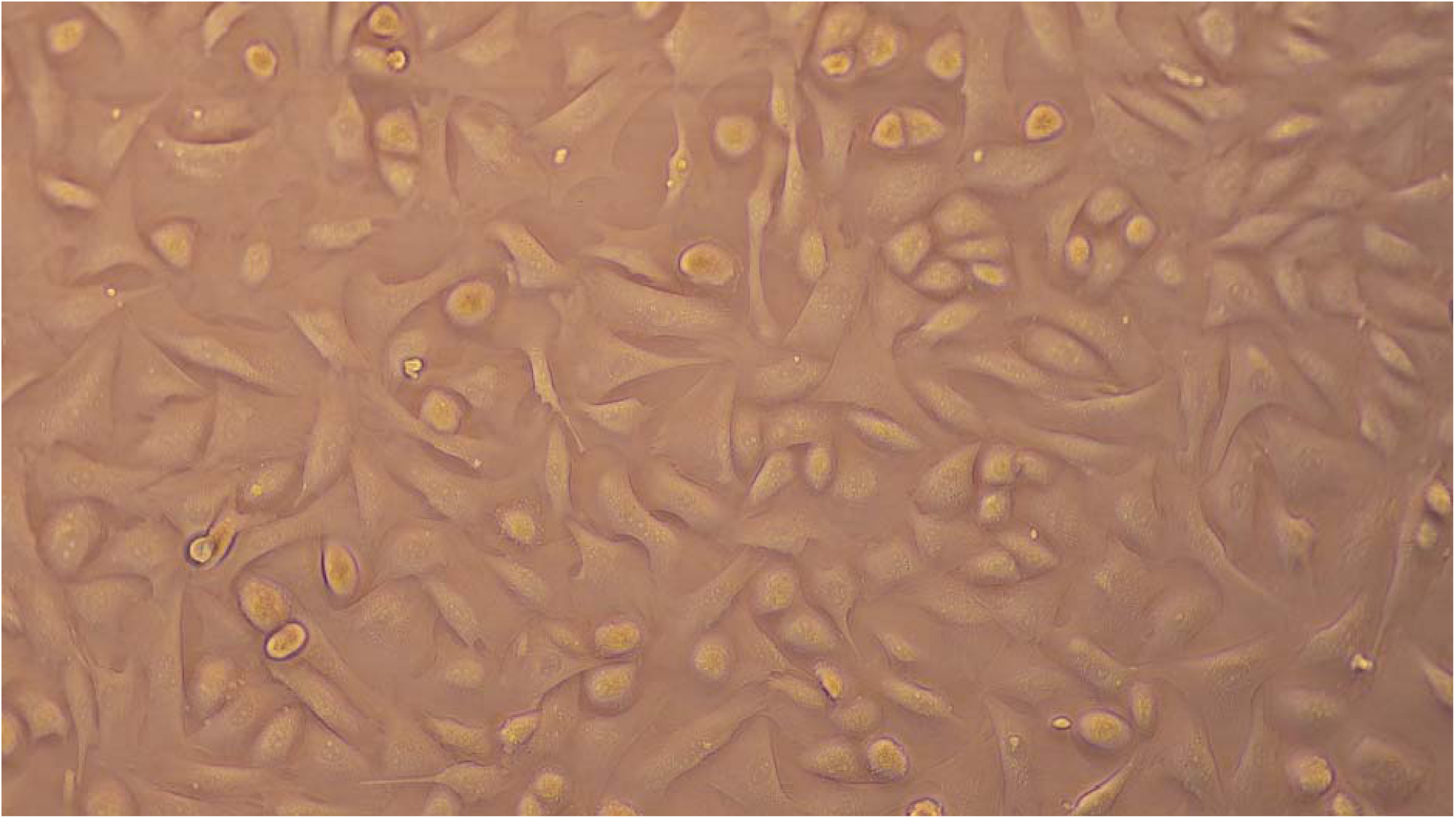
The heart cell line passage 30 (three hrs post incubation). The first generation of cells after the spontaneous cell fusion in passage 29 resulted after trypsinization of the 72 hrs multicellular filamentous growth in passage 29, bright field X200.

**Fig 8.**
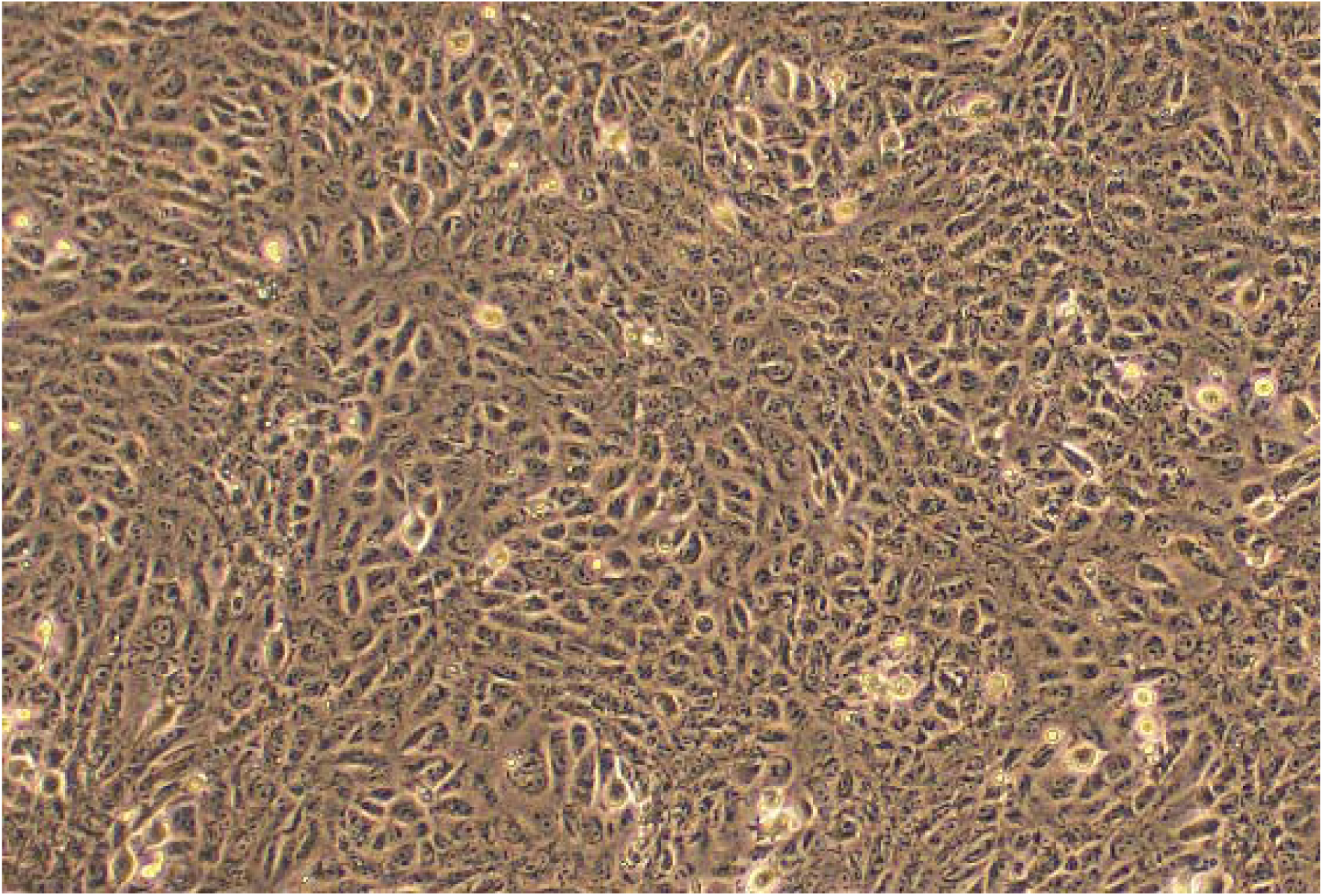
Morphology of the heart cell line passage 33 after 48 hrs incubation (this morphology remained constant in the subsequent subcultures till passage 140, phase contrast X100.

Although the final cell line morphology was epithelial-like cell morphology in fact it consisted of two morphological types fibroblast-like cells and epithelial-like cells similar in morphology to parent cells, but this distinction could only be detected during the first 3-5 hours of culture incubation, after which the cell line appeared as one cell morphology.

When we examined many fields of passage 30 culture, points of cell fusion surrounded by a matrix of the second generation growing progeny cells were detected. These fused cells represented cells that had failed to fuse in passage 29 due to physical distancing. Such cells possessed exceptionally large nuclei with prominent nucleoli. The events following cell fusion proceeded as described before; moreover, the nuclear fusion was preceded by clear disintegration of the nuclear membranes of the fusing nuclei and obvious blending of the genetic material of the two nuclei. In addition, the nuclear reaction was accompanied with very brilliant glow indicative of the high energy provided by the mitochondria to complete the fusion reaction (Fig. 9 A-F).

**Fig 9.**
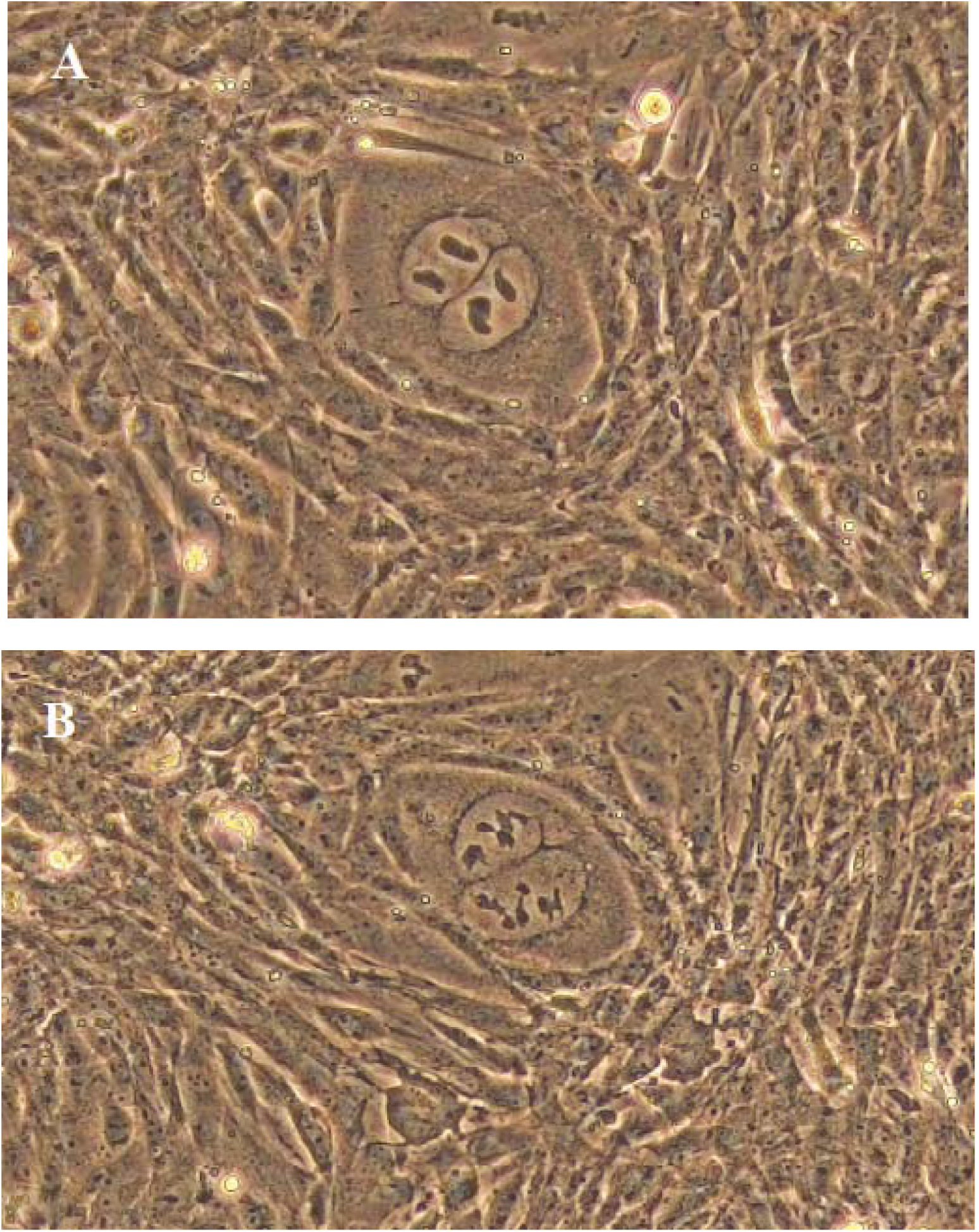
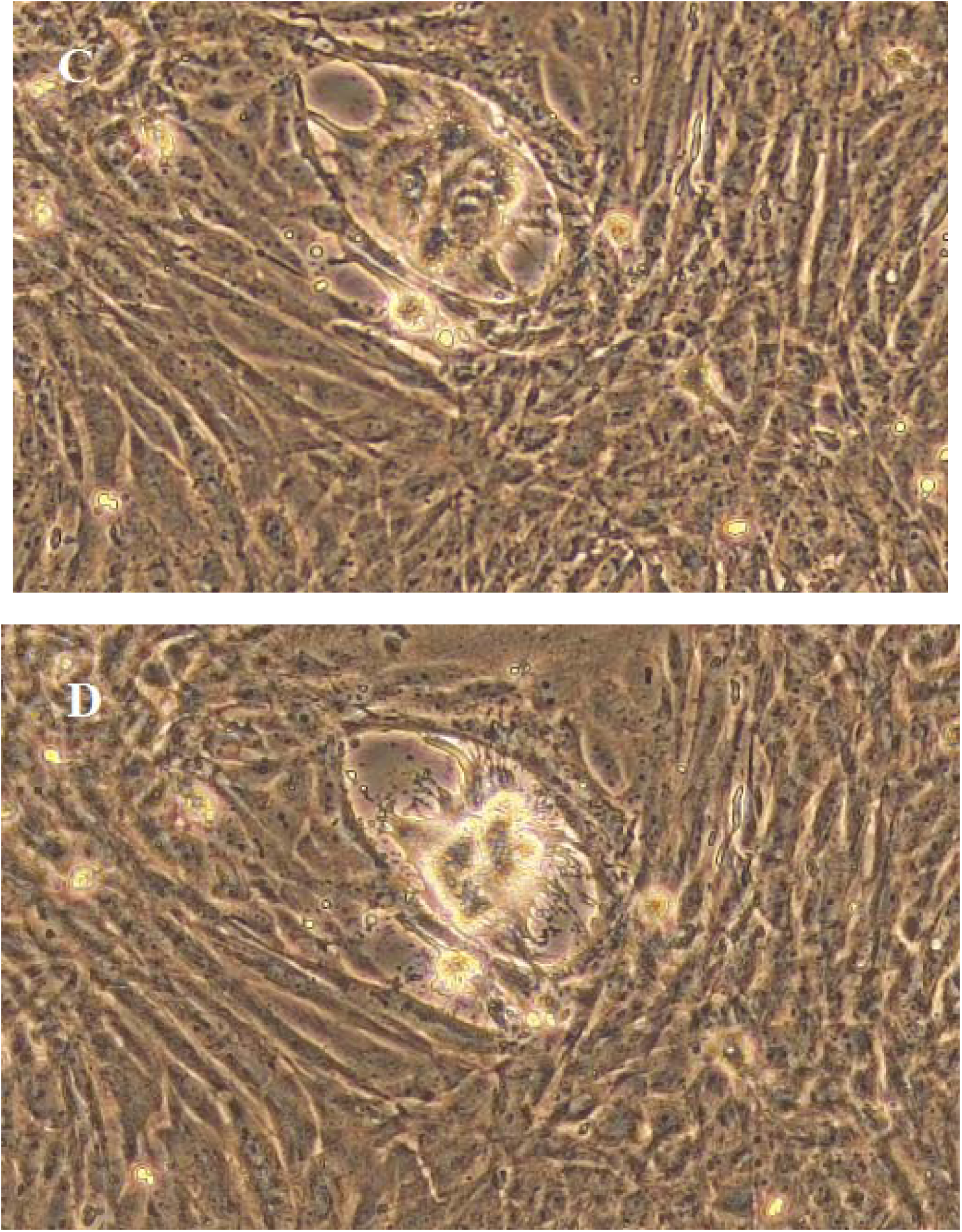
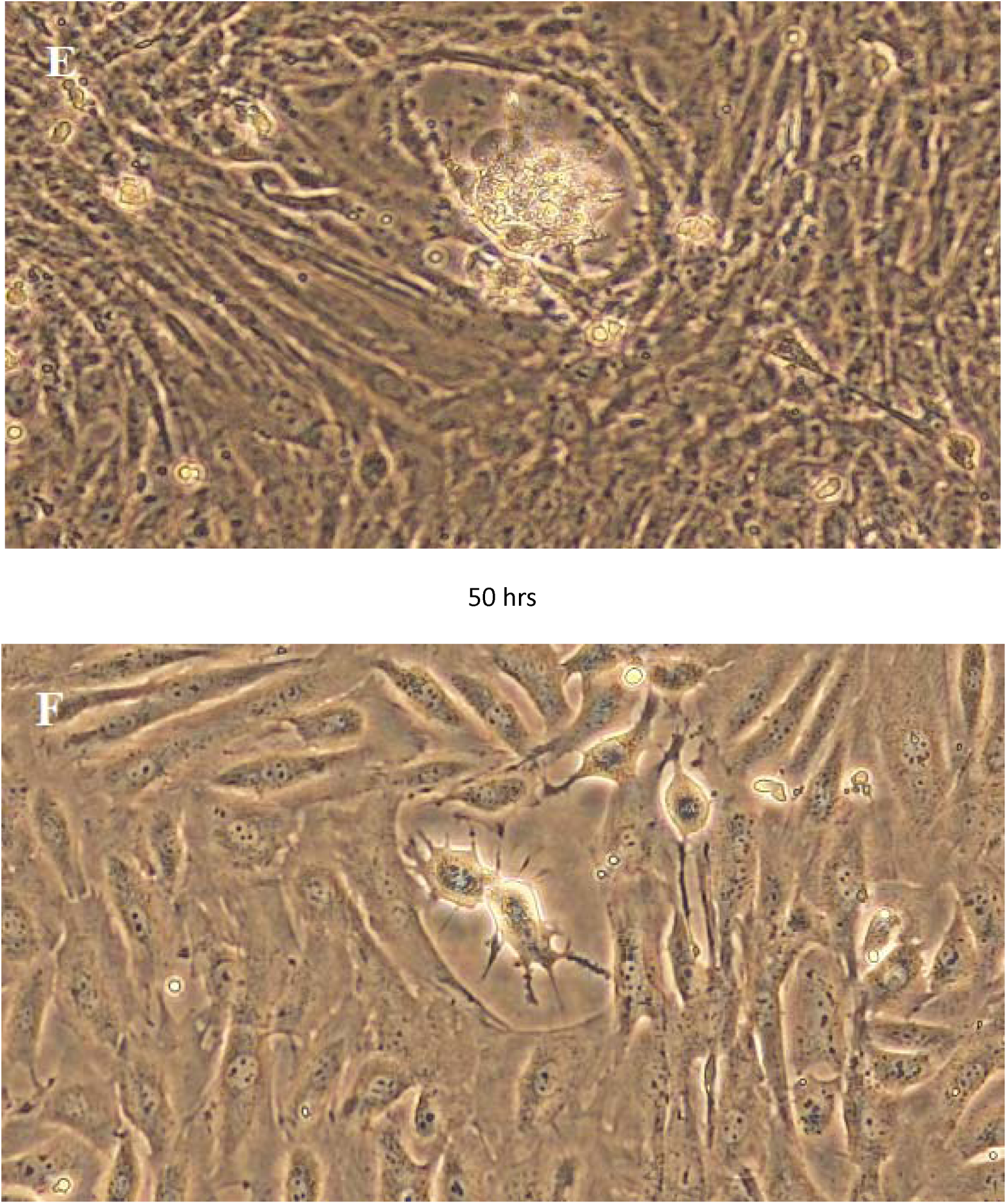
A point of spontaneous cell fusion in passage 30. a) 24 hrs post incubation culture showing cell fusion and coupling of nuclei with accumulation of mitochondria around them, b) 46 hrs post incubation, disintegration of the nuclear membranes of the two nuclei and rearrangement of their genetic material, c) 48 hrs post incubation, complete mix up of the genetic material of the two cells, d) 49 hrs post incubation, nuclear fusion reaction with massive liberation of energy (burning of the genetic material), e) 50 hrs post incubation, cooling down of the nuclear fusion reaction and reappearance of the genetic material for the developing progeny cells, and f) two nuclei separating at the end of the nuclear fusion reaction (Fig. (F) was photographed from a different a different culture), phase contrast X200.

In general, during the study of the spontaneous cell fusion between the two heart cell phenotypes, we observed that the cell fusion process was photosensitive and prolonged exposure of a fixed field during documentation by photography resulted in delay and sometimes abortion of the cell fusion process in the specific field under examination.

### Growth Curve and population doubling time

A characteristic growth curve of cells after transformation established using a three day old culture of passage 36. The curve showed no lag phase and the highest growth rate was in the first 24 hours and the log phase continued for 10 days. Transformed cells had a population doubling time of 14.5 hours compared to 25.8 hours for cells before transformation. The growth curve of the heart cell line passage 36 is shown in S1 Fig.

### Storage of the cell line at 37° C

The FOH-SA cell line was successfully subcultured after six months storage at 37° C, however, trypsinization of the culture required 30 minutes rather than the normal 5 minutes and the whole monolayer came out as one sheet which was broken down after rigorous shaking and pipetting. During the storage period the cell monolayer showed no signs of degeneration, however, it was noteworthy that numerous islets could be detected in the monolayer in which the cells again acquired the ability to multiply in multicellular filaments, similar to the filamentous cell morphology reported in the transitional passages 29-32 (S2 Fig.). The cells after storage remained sensitive to sheep pox virus and its CPE remained similar in pattern to the original cells.

### Cell line authentication

Authentication of the ovine heart cell line at ECACC revealed that the mitochondrial DNA barcode of passage 22 cells matched 99% to *Ovis aries* (top 100 results on BLAST). Result of mycoplasma PCR analysis at ECACC documented that the heart cell line was mycoplasma free.

### SNP genotyping

The OvineSNP50 Beadchip map contained 54241 ovine SNPs spanning all the 26 autosomal chromosomes, the sex chromosomes and the mitochondria. The genotyping data of the DNA samples of passage 22 and 47 generated by the Infinium assay of the bead chip on the iScan® system were analyzed by the software Illumina GenomeStudio version 2.0.2 and GenCall Version 7.0.0 with low GenCall Score Cutoff equal to 0.050.

A total of 50653 SNPs were successfully called (representing 93.4% of the total number of SNPs on the bead chip) in DNA sample of passage 22 cells. The frequency the genotype of the SNP alleles was 0.65 homozygous and 0.35 heterozygous.

45774 SNPs were called in the DNA sample of passage 47 which represented 84.4% of the total number of SNPs on the bead chip. The frequency of genotype of the alleles was 0.96 and 0.04 for homozygosity and heterozygosity respectively.

The summary of the total number of SNPs called, the genotypes of the SNPs, the minor allele frequency, and the 50% GenCall score of each of the DNA samples is shown in table 1.

**Table 1:** OvineSNP50 BeadChip genotyping of fetal ovine heart cell line.

| DNA<br>sample | # SNPs<br>called | A/A<br>Freq | A/B<br>Freq | B/B<br>Freq | MAF | 50%<br>GC<br>Score |
| --- | --- | --- | --- | --- | --- | --- |
| Passage 22 | 50653 | 0.30 | 0.35 | 0.35 | 0.48 | 0.903 |
| Passage 47 | 45774 | 0.95 | 0.04 | 0.01 | 0.03 | 0.875 |

Cell line before cell fusion (passage 22), Cell line after cell usion (passage 47), the total number of SNPs in each sample with GenCall score above the no-call threshold (#SNPs called), the number of A/A SNP genotype calls divided by the total number of calls (A/A Freq), the number of A/B SNP genotype calls divided by the total number of calls (A/B Freq), the number of B/B SNP genotype calls divided by the total number of calls (B/B Freq), Minor allele frequency (MAF), 50% GenCall Score (50% GC score).

4879 SNPs which were called in passage 22 were not called in passage 47 sample. The no-all SNPs were distributed throughout the chromosomes; the highest numbers were reported in chromosomes 3, 2, 1, 5, and 9. The no-call SNPs occurred singly and in succession of two, three, four, and five SNPs. The genes spanning the three, four, and five in succession no-call SNPs were identified by inserting the location of each set of SNPs in ENSEMBL (S1 Table).

The genetic conversion was also documented in the mitochondrial SNPs *mt.7729*, *CytB_1505.1*, and *CytB_1745.1*. The alleles of these SNPs were 100% BB homozygous in passage 22 cells (BT) became 67% AA and 33% BB homozygous in passage 47 cells (AT).

The single SNP in the Y chromosome (*oY1.1,* index 37199 in the SNP map) was neither called in the cell line before transformation nor called in the cells after transformation; accordingly we concluded that the cell line was established from a female ovine fetus since the sex of the fetus was not checked during the preparation of primary cultures.

### Sequencing of cytochrome b gene

Sequencing of cytochrome b gene of the heart cell line passage 26 and 59 produced sequences of 1078 bp and 1061 bp respectively. After quality control and trimming the length of *cytb* gene was 834 bp for passage 26 cells and 858 bp for passage 59 cells. Blastn analysis of *cytb* of passage 26 cells revealed 99.28% identity to 100 BLAST Hits of *Ovis aries* complete mitogenome. Pairwise alignment of the partial sequences of *cytb* genes of the two cell passages showed 93.8 % similarity with 6.2% gaps (S3 Fig.). Blastn analysis of the partial sequence of *cytb* gene of passage 59 cells showed 99.88% identity to the top 100 Blast Hits of *Ovis aries* mitogenome and this sequence was deposited with GenBank database (GenBank accession: ON811684).

### Sequencing of the control region, the *tRNA-Phe* and *12S rRNA* genes

The partial sequencing of the control, the *tRNA-Phe* gene and the 12S *rRNA* genes of the heart cell line mitogenome resulted in a sequence of 1088 bp for passage 26 cells and a 1083 bp for passage 59 cells, however, after trimming and quality control, the size of sequences was 828 bp for passage 26 cells and was only 305 bp for passage 59 cells. The 100 top Blast Hits for passage 26 partial sequence revealed 98.67-99.88% identity to *Ovis aries* control region; *tRNA-Phe* and 12S *rRNA* mitogenome genes. Pairwise sequence alignment of the 828 bp sequence of passage 26 and 305 bp sequence of passage 59 revealed only 25.6% similarity with 67.9 % gaps (S4 Fig.).

### Susceptibility to viruses

The FOH-SA cell line was found highly susceptible and permissive to all tested viruses. The CPE of RVF virus was detected within 48 hours in the form of cell elongation, syncytia formation, and disruption of the monolayer. RVF virus suspension was harvested after four days and the TCID_50_ was 10 ^7.1^/ ml. The CPE PPR virus developed within 72 hrs in form of cell rounding, syncytia formation and the monolayer remained adherent to surface. The virus was harvested after five days and the TCID_50_ of the harvest reached 10^7.3^/ml. LSD virus showed CPE similar to that of PPR, however, it dismantled the monolayer sheet. The tire of LSD virus harvest was 10^7.5^ TCID_50_/ ml after five days incubation. The CPE of camel pox virus was detected after 48 hours in the form of rounding of cells, large syncytia, and detachment of the monolayer (10^7.5^ TCID_50_/ml after 3 days). The CPE of sheep pox virus after 48 hrs and the titre of the sheep pox virus was harvested after 5 days (10^7.5^ TCID_50_/ ml). CPE of sheeppox, camelpox, and PPR viruses is shown in S5 Fig.

### Production sheep pox vaccine

The cell line was used since its establishment in 2013 mainly in production of sheep pox and camelpox vaccines. Over the years production was carried out in cell line passages ranging from 38 to 115 and satisfactory viral suspensions with titres of 10^7.1^TCID_50_/ ml to 10^7.5^ TCID_50_/ ml for both vaccines were obtained.

### ATCC Cell line deposition

FOH-SA cell line passed all the safety tests performed by FADDL including freedom from FMD, PPR, sheep and camel pox viruses, then it was shipped to ATCC where it was processed in accordance with Budapest Treaty and the relevant international Budapest Treaty forms were issued and the cell line was deposited under patent accession number (PTA: 125751).

## DISCUSSION

Spontaneous immortalization of mammalian cells is rare happening event where spontaneous up regulation of telomerase reverse transcriptase enzyme or the low expression p15/p16/Rb allow cells to escape replication senescence and become immortal [24,25,26].

Although numerous animal cells lines have been spontaneously immortalized by serial passage, the extract mechanisms causing cell immortalization remained unexplained. For instance the mechanisms that led to the establishment of the African green monkey cell line (VERO), which was derived by serial passaging of normal kidney cells, remained unexplained though the cell line was established decades ago.

The transformation event documented in this study passed undetected during the initial serial passaging to establish an immortal heart cell line, however, the sufficient store of before transformation cells allowed us to repeatedly derive the cells through passage 29 which enabled us to precisely observe and explain of the events that led to cell transformation, otherwise the transformation event reported in this study would have remained unexplained. Ogle *et al.* [27] reviewed that the origin of a cell as the product of fusion event can be difficult or impossible to deduce.

In this study we established an immortal fetal ovine heart cell line by serial passaging and we described unprecedented method of cell immortalization in which two heart cell phenotypes spontaneously fused and then segregated into two progeny cells that have different morphology and smaller in size compared to parent cells, have an increased proliferation potential, exhibited a transitional phase of filamentous growth, immortal, and susceptible to numerous animal viruses.

Fusion of cells of the same or of different type and fusion of their nuclei was found to occur *in vivo* yielding hybrid cells known heterokaryons (multinuclear) or synkaryons (mononuclear). In synkaryons resorting and recombination of chromosomal DNA could occur and the mononuclear daughter cells resulting from the hybrid cells were found to express all the genetic material of the parental cells [28, 29].

The spontaneous cell fusion between the fibroblast-like and epithelial cell-like heart cell phenotypes was repeatedly documented at passage 29 (cells were subcultured every 3-4 days) and was not detected at any earlier passage and this could mean that these cell types become mature and competent for fusion at this specific age, 29 passages, possibly by expressing the necessary cell fusion proteins and having mature mitochondria capable of providing the energy necessary for the nuclear fusion reaction. Recent work has identified fusion proteins (fusogens) as indispensible molecules in the fusion process between mammalian myoblast cells [28, 30], similar fusogens might have mediated the fusion of the heart cells documented in this study.

In this study we observed two types of spontaneous cell fusion; adjacent fusion occurred when the two phenotypes existed in neighborhood and was detected within 3-5 hours after incubation of passage 29 cells, and a second type of cell fusion occurred when the cell types were distantly placed from each other and here the fibroblast-like cells protruded projections reaching the epithelial-like cells and eventually a tubule was established connecting the two cells. Shortly after cell fusion, coupling of the nuclei of the fused cells occurred which triggered the migration of mitochondrial population of the two cells to the site of nuclear coupling. The force driving the two nuclei to a coupling site mid the tubular junction between the two cells could not be explained in this study. The mitochondrial migration was found to be crucial for initiating the nuclear fusion process and it was only triggered when all the mitochondria of the fusing cells accumulated at the site of coupled nuclei.

Cell fusion *in vitro* was induced by treating human lymphocytes with polyethylene glycol [31]. A second approach for cell fusion *in vitro*, known as electrofusion, was experimented by applying electric impulses to cells suspended in an appropriate electrofusion buffer [32, 33]. Both methods resulted in hybrid cells with two or three nuclei, however, no nuclear fusion between the nuclei of hybrid cells were demonstrated in these studies, hence the spontaneous cell fusion and nuclear fusion described in this study was a novel unprecedented phenomenon in cell biology.

The spontaneous cell fusion, described in this study, between the two heart cell phenotypes at passage 29 could be viewed as a type of cell mating (sexual reproduction), which is followed by a form of cell multiplication that could considered as asexual reproduction in which each progeny cell grew into multiple cells that remained connected to each other forming filaments. This pattern of cell multiplication was documented in the transitional passages 29, 30, 31, and 32 and could described as resembling septate hayphae resulting from asexual reproduction in fungi. An early report described *in vitro* mating between two clonal lines of mouse fibroblasts that resulted in hybrid cells with evidence of segregation after mating, however, research in cell mating was discontinued and no other reports are available [34].

The OvineSNP50 BeadChip has been used in many applications which included genome-wide selection, comparative genetic studies, and breed characterization for evaluating biodiversity [35, 36, 37]. It was validated by Illumina for genotyping of more than 3000 samples from diverse *Ovis aries* domestic and wild breeds [19]. In the current study we used the BeadChip to compare the genotype of the cell line before and after cell fusion. SNP genotyping of DNA of heart cell line passage 22 revealed 50653 polymorphic loci representing 93.4% of the SNPs on the bead chip which was higher than the highest number of polymorphic loci reported in all domestic ovine breeds (*Rasa aragonesa* breed of sheep exhibited the highest polymorphic loci of 48676) during the validation of the bead chip by Illumina, though the *Harri* sheep breed, the sheep breed from which the heart cell line was established, was not included in the buildup of the bead chip.

SNP genotyping of passage 47 cells showed a 0.95 A/A and 0.01 B/B SNP genotype frequency compared to 0.30 A/A and 0.35 B/B genotype frequencies in DNA samples of passage 22, while the frequency for heterozygosity was 0.35 and 0.04 for passage 22 and 47 respectively. These results indicated that the spontaneous cell fusion resulted in a large scale genetic conversion in progeny cells involving about 65% of the alleles resulting in 96% homozygosity. Osada et al. [38], identifying the single nucleotide variants (SNV) of the African green monkey (*Chlorocebus sabaeus*) cell line (VERO) nuclear genome, reported 87.7% homozygosity and they attributed the high homozygosity to evolutionary divergence between the genus *Macaca* and *Chlorocebus* approximately 8-12 million years ago.

The minor allele frequency (MAF) of the heart cell line before cell fusion was 0.48 after cell fusion was 0.03 The MAF of 0.03 for FOH-SA cell line after cell fusion indicated that FOH-SA cell line was a rare variant possessing high level of rare alleles, which is consistent with fact that spontaneous cell fusion in animal cells is a rare event.

The ovine mitochondrial *cytb* gene and a fragment of the ovine mitogenome containing a control region have been used to haplotype sheep breeds and to study genetic diversity among and between sheep breeds [20, 21, 39]. In this study we partially sequenced the *cytb* gene and the mitochondrial fragment containing the control region and *tRNA-Phe* and 12S *rRNA* genes of the heart cell line to explore whether the spontaneous cell fusion has led to mutational events in the mitogenome of progeny cells. Pairwise sequence alignment of the partial sequences of the cell line passage 26 and passage 59 revealed 54 gaps caused by insertion 35 bp in the 5’ end and 15 deletion in 3’ end of the partial sequence of cytb of passage 59 cells. These insertion/ deletion events in the cytb of passage 59 might have caused the change in the genotypes of *Cytb* gene SNPs as it was documented by the Ovine bead chip genotyping of passage 47 of the FOH-SA cell line.

25.6% was the similarity between the mitogenome segments spanning the control region, the *tRNA-Phe* and 12S *rRNA* genes of heart cell line passages 26 and 59. The low similarity was caused by 583 bp gaps resulting from major deletion events in the segment of passage 59 cells mitogenome. The results of partial sequencing of the mitogenome of heart cell line substantiated the results of SNP genotyping for the occurrence of large scale genetic conversion in the progeny cells following the spontaneous cell fusion. Osada *et al.* [38] sequenced the complete mitogenome of Vero cells and phylogenetically it was so diverse from *Chlorocebus sabaeus* (the species from which it was established) to the extent that the Vero cell clustered as a separate species.

We believe that the events that led to FOH-SA cell line development and Vero cell line establishment were analogous i.e. both cell lines evolved by spontaneous cell fusion. This reality was ascertained first, by the fact that a three hour culture of FOH-SA cells and that of VERO cell line looked morphologically identical both have endothelial-like morphology (S6 Fig.). Second, The endothelial-like morphology of a three-hour culture of both cell lines is due to the fact that both cell lines consisted of two cell types, one type descending from a fibroblast-like cell usually showing rudimentary projections few hours after incubation that soon disappear with growth, and an epithelial-like cell, a morphological feature indicative of the ancestors of the two cell lines and comparing a three hour culture of Vero ells with heart cell line passage 22 (Before transformation cells) strikingly demonstrated this fact (S7 Fig.). Third, the large scale genetic conversion in both cell lines resulting in high level of homozygosity in both cell lines a fact that could only be ascribed to spontaneous cell fusion. Fourth, the major mutational events detected in the mitogenome of both cell lines. Future comparative studies of the mitogenomes of the FOH-SA and Vero cell lines will confirm the existence in the animal body of cell phenotypes that could spontaneously fuse giving rise to progeny cells that are genetically highly divergent from the parent cells.

The cell line was found highly permissive to rift valley fever, PPR, lumpy skin disease, camel pox and sheep pox viruses; hence, it could be used for isolation of these viruses as well as development and production of vaccines. Immortalization of cells was found to be associated with silencing or deletion of the interferon (IFN) gene cluster [40] and the sensitivity of Vero cells to numerous viruses was also ascribed to deletion of the IFN gene cluster [38]. Similarly, the sensitivity of FOH-SA to these viruses might be attributed to deletion of IFN gene cluster; a future genome landscape of the cell line would disclose this fact. The severe acute respiratory syndrome coronaviruses 2 and 1 (SARS-CoV-2, SAR-CoV-1) utilize the angiotensin converting enzyme 2 (ACE2) as a receptor for cell entry and the ACE2 receptor is abundantly expressed on cardiocytes, cardiofibroblasts and coronary endothelium [41, 42], hence, the FOH-SA cell line being derived from heart tissue would be a good candidate for propagation of SARS-CoV-2 and SARS-CoV-1 viruses.

A unique property of the FOH-SA cell line was the possibility of storing cultured cells at 37° C for months provided that the growth medium was changed every three weeks. This property could be exploited as a short storage mechanism avoiding the cumbersome and risky cryopreservation of cells when they are needed within months and also this property could be useful in propagation of slow growing viruses. Shipment of this cell line could be done using cultures which is more practical and economic than shipping on dry ice or liquid nitrogen. The capacity of FOH-SA cells stored at 37° C to revert to multicellular filamentous growth similar to the filamentous growth documented in passages 29 to 32 would be a characteristic differential property for this cell line.

## Acknowledgements

We would like to acknowledge Mohamed M. Alfuhaid the Director General of the General Administration of Laboratories for his continuous support, encouragement, and enthusiasm. We greatly appreciate the close follow up and the unlimited support of Dr. Khaled S. Abuhimed, Director of Veterinary Vaccine Production and Evaluation Centre. The help of Dr. Hussain Al-Ghadeer and Dr. Elgazali Guma, Veterinary Diagnostic Laboratory, MEWA, Riyadh, in preparing DNA samples is greatly appreciated.

## Author contributions

Khalid. M. Suleiman: Designed and investigated the study, analyzed the results, and prepared the manuscript.

Mutaib M. Aljulidan: Participated in preparation of primary cell culture, passaging, and cryopreservation.

Gamal eldin M. Hussein: Collaborated in preparing primary cell culture and testing the sensitivity of the cell line to viruses.

Habib N. Alkhalaf: Participated in designing the study and supervised it.

## REFERENCES

1. Eibl D, Eibl R, Poertner. Cell and Tissue Reaction Engineering: Principles and Practice, ® Springer Berlin Heidelberg 2008.

2. Swain PK, Nayak SK, Mishon SS. Basic technique and limitations in establishing cell cultures: A mini review. Ad An Vet Sci. 2014; 2: 1-10.

3. Verma A, Verma M, Singh A. Animal tissue culture principles and applications. Animal Biotechnology (Second Edition), Academic Press, 2020: 269-293.

4. Lloki Assanga SB, Gil-Salido AA, Lewis Lujan LM, Rosas-Durazo A, Acosta-Silva AL, Rivera-Castaneda EG, Rubio-Pino JL. Cell growth curves of different cell lines and their relationship with biological activities. Inter J Biotech Mol Bio Res. 2013; 4: 60–70

5. Suleiman KM, Boehnel H, Babiker SH, Zaki AZSA. Cytotoxic activity of fermenter culture supernatants of *Corynebacteriumpseudotuberculosis*. J Anim Vet Ad. 2006; 5: 939–942.

6. Soice E, Johnston J. Immortalizing Cells for Human Consumption. Int J Mol Sci. 2021; 22: 11660.

7. Hayflick L, Moorhead PS. The serial cultivation of human diploid cell strains. Exp Cell Res. 1961; 25: 585–621.

8. Hayflick L. The cell biology of aging. J Investig Dermatol. 1979; 73: 8–14.

9. Kuilman T, Michaloglou C, Mooi WJ, Peeper DS. The essence of senescence. Genes Dev. 2010; 24: 2463–2479.

10. Alessio N, Squillaro T, Cipollaro M, Bagella L, Giordano A, Galderisi U. The BRG1 ATPase of chromatin remodeling complexes is involved in modulation of mesenchymal stem cell senescence through RB-P53 pathways. Oncogene 2010; 29: 5452–63.

11. Kumari R, Jat P. Mechanisms of Cellular Senescence: Cell Cycle Arrest and Senescence Associated Secretory Phenotype. Front Cell Dev Biol. 2021; 9:645–593.

12. Friedman AL, Tainsky MA. Critical pathways in cellular senescence and immortalization revealed by gene expression profiling.Oncogene. 2008; 27(46): 5975–5987.

13. Shay JW, and Wright WE. Telomeres and telomerase: Three decades of progress. Nat Rev Genet. 2019; 20: 299–309.

14. Madin Sh, Darby NB Jr. Established kidney cell lines of normal adult bovine and ovine origin. Proc Soc Exp Biol Med. 1956; 98: 574–576.

15. Matsuura K, Inoshima Y, Kameyama K, Murakami K. Establishment of a novel ovine kidney cell line for isolation and propagation of viruses infecting domestic cloven-hoofed animal species. In Vitro Cell De Biol Anim. 2011; 47:459–463.

16. History and Characterization of the Vero Cell Line- A Report prepared by CDR Rebecca Sheets, Ph.D., USPHS CBER/OVRR/DVRPA/VVB for Vaccines and Related Biological Products Advisory Committee Meeting to be held on May 12, 2000 OPEN SESSION.

17. Davis JM. Basic Cell Culture, a Practical Approach. © Oxford University Press; 1994.

18. Herbert PD, Cywinska A, Ball SL, de Waard JR. Biological identifications through DNA barcodes. Proc Biol Sci. 2003; 270: 313–321.

19. Meadows JR, Li K, Kantanen J, Tapio M, Sipos W, Pardeshi V, Gupta V, Calvo JH, Whan V, Norris B, Kijas JW. Mitochondrial sequence reveals high levels of gene flow between sheep breeds from Asia and Europe. J Hered. 2005; 96: 494– 501.

20. Hiendleder S, Lewalski H, Wassmuth R, and Janke A. The complete mitochondrial DNA sequence of the domestic sheep (*Ovis aries*) and comparison with the other major ovine haplotype. J Mol Evol. 1998; 47: 441–448.

21. Karber G. Beitrag zur kollektiven Behandlung pharmakolgischer Reihenversuche. Archiv f experiment Path u Pharmakol. 1931; 162: 480–483.

22. OvineSNP50 Genotyping BeadChip-Illumina. https://www.illumina.com/documents/products/datasheets/datasheet_ovinesnp50.pdf

23. Madeira F, Pearce M, Tivey ARN, Basutkar P, Lee J, Edbali O, Madhusoodanan N, Kolesnikov A, Lopez R. Search and sequence analysis tools services from EMBL-EBI in 2022. Nucleic Acids Research, 12 Apr 2022,:gkac240 DOI: 10.1093/nar/gkac240 PMID: 35412617.

24. Shay JW, Wright WE. Senescence and immortalization: role of telomeres and telomerase. Carcinogenesis 2005; 26:867–874.

25. Lundberg AS, Hahn WC, Gupta P, Weinberg RA (2000). Genes involved in senescence and immortalization. Curr Opin Cell Biol. 2000; 12:705–9.

26. Visser TB, van Veen EA, van Meerloo J, Braakhuis RH, Steenbergen RD, Brakenhoof RH. Immortalization of oral kratinocytes by functional inactivation of the p53 and pRB pathways. Int J Cancer. 2011; 128: 1596–1605.

27. Ogle BM, Cascahlo M, Platt JL. Biological Implication of cell fusion. Nat Rev Mol Cell Biol. 2005; 6: 567–75.

28. Zhang H, Ma H, Yang X, Fan L, Tian S, Niu R, Yan M, Zheng M and Zhang S. Cell Fusion-Related Proteins and Signaling Pathways, and Their Roles in the Development and Progression of Cancer. Front. Cell Dev. Biol. 2022; 9:809–668. doi: 10.3389/fcell.2021.809668.

29. Alvarez-Dolado M. Cell fusion: biological perspective and potential for regenerative medicine. Front Biosci. 2007; 12: 1–12.

30. Bi P, Ramirez-Martinez A, Li H, Cannavino J, McAnally JR, Shelton JM, Sanchez-Ortiz E, Bassel-Duby R, Olson EN. Control of muscle formation by the fusogenic micropeptide myomixer. Science. 2017; 356: 323–327.

31. Pedrazzoli F, Chrysantzas I, Dezzani L., et al. Cell fusion in tumor progression: the isolation of cell fusion products by physical methods. Cancer Cell Internal. 2011; 11: 32

32. Kanduser M, Usaj M. Cell electrofusion: past and future perspectives for antibody production and cancer cell vaccines. Expert Opin. Drug Deliv. 2014; 11: 1–14

33. Ramos C, Teissie J. Electrofusion: a biological modification of cell membrane and mechanism in exocytosis. Biochimie. 2000; 82: 511–518

34. Littlefield JW. Selection of hybrids from mating of fibroblasts in vitro and their presumed recombinants. Science. 1964; 145: 709–710

35. Sandenbergh L, Cloete SWP, Roodt-Wilding R, Snyman MA, Bester-van der Merwe AE. Evaluation of the OvineSNP50 chip for use in four South African sheep breeds. S. Afr. J. anim. 2016; 46: 89–93.

36. Ilori BM, Rosen BD, Sonstegard TS, Bankole OM, Durosaro SO, Hanotte O. Assessment of OvineSNP50 in Nigerian and Kenyan sheep populations (2018). Nig. J. Biotech. 2018; 35 (2): 176–183.

37. Cao Y, Song X, Shan H, Jiang J, Xiong P, Wu J, Shi F and Jiang Y Genome-Wide Association Study of Body Weights in Hu Sheep and Population Verification of Related Single-Nucleotide Polymorphisms. Front. Genet. 2020;11:588.

38. Osada N, Kohara A, Yamaji T, Hirayama N, Kasai F, Sekizuka T, Kuroda M, and Hanada K. The genome landscape of the African green monkey kidney-derived Vero cell line. DNA Res. 2014; 21: 673–683.

39. Othman EO,, Germot A,, Khodary MG, Petit D, Maftah A (2018). Cytochrome b diversity and phylogeny of six Egyptian sheep breeds. Annual Research & Review in Biology 22:1–11.

40. Fridman AL, Tainsky MA. Critical pathways in cellular senescence and immortalization revealed by gene expression profiling. Oncogene. 2008; 27: 5975–5987.

41. Tai W, He L, Zhang X., et al. Characterization of the receptor binding domain of 2019 novel coronavirus: implication for development of RBD proteins as a viral attachment inhibition and vaccine. Cell Mol Immunol. 2020; 17: 613–620.

42. Patel VB, Zhong JC, Grant MB, Oudit GY. Role of the ACE2/ Angiotensin 1-7 Axis of the Rein-Angiotensin system in heart Failure. Circulation Res. 2016; 118: 1313–1326.

