## Supplemental files for "Establishment of a novel fetal ovine heart cell line by spontaneous cell fusion"

Habib N. Alkhalaf^a^

a. Veterinary Vaccine Production and Evaluation Centre, Ministry of Environment, Water, and Agriculture, Riyadh, Saudi Arabia.

**­­­­­­­­­­­­­­_________**

*Corresponding author: address: Veterinary Vaccine Production and Evaluation Centre, Ministry of Environment, Water, and Agriculture, Riyadh, 11454, KSA.

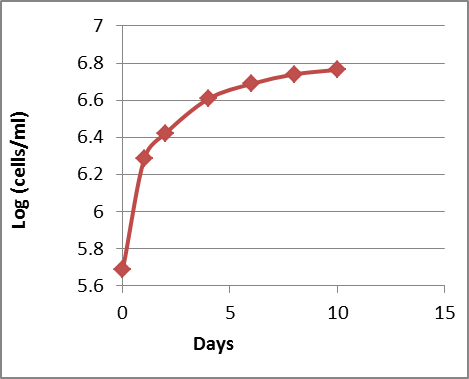

**S1Fig**.Growth curve of FOH-SA cell line passage 36.

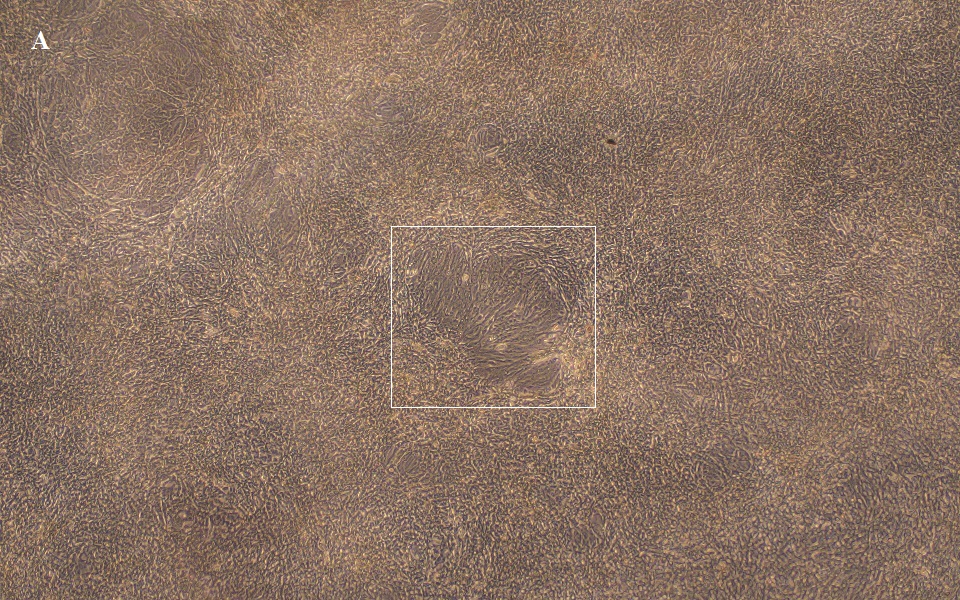

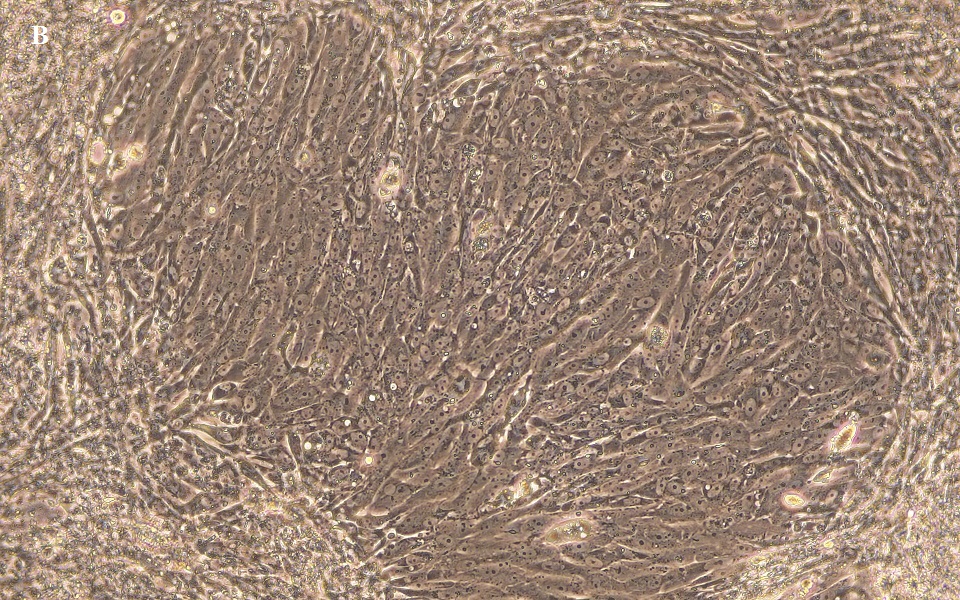

**S2 Fig**. a) the monolayer of FOH-SA cell line passage 59 after 6 month storage at 37 °C, phase contrast X100, b) the rectangular central area magnified X400 to show the cell growing into multicellular filaments.

```
########################################
# Program: needle
# Rundate: Sun 19 Jun 2022 19:49:06
# Commandline: needle
#    -auto
#    -stdout
#    -asequence emboss_needle-E20220619-194905-0048-38661723-p1m.aupfile
#    -bsequence emboss_needle-E20220619-194905-0048-38661723-p1m.bupfile
#    -datafile EDNAFULL
#    -gapopen 10.0
#    -gapextend 0.5
#    -endopen 10.0
#    -endextend 0.5
#    -aformat3 pair
#    -snucleotide1
#    -snucleotide2
# Align_format: pair
# Report_file: stdout
########################################

#=======================================
#
# Aligned_sequences: 2
# 1: A8_cytb-F1_20220424_SAMPLE1-MGCOMP-28572437
# 2: C8_cytb-F2_20220424_SAMPLEA-36136530
# Matrix: EDNAFULL
# Gap_penalty: 10.0
# Extend_penalty: 0.5
#
# Length: 873
# Identity:     817/873 (93.6%)
# Similarity:   819/873 (93.8%)
# Gaps:          54/873 (6.2%)
# Score: 4056.5
#
#
#=======================================

A8_cytb-F1_20      1 -----------------------------------TAATGATCAACATCC     15
                                                        |||||||||||||||
C8_cytb-F2_20      1 GAAAAACCATCGTTGTCATTCAACTATAARAACACTAATGATCAACATCC     50

A8_cytb-F1_20     16 GAAAAACCCAYYCACTAATAAAAATTGTAAACAACGCATTCATTGATCTC     65
                     ||||||||||..||||||||||||||||||||||||||||||||||||||
C8_cytb-F2_20     51 GAAAAACCCACCCACTAATAAAAATTGTAAACAACGCATTCATTGATCTC    100

A8_cytb-F1_20     66 CCAGCTCCATCAAATATTTCATCATGATGAAACTTTGGCTCTCTCCTAGG    115
                     ||||||||||||||||||||||||||||||||||||||||||||||||||
C8_cytb-F2_20    101 CCAGCTCCATCAAATATTTCATCATGATGAAACTTTGGCTCTCTCCTAGG    150

A8_cytb-F1_20    116 CATTTGCTTAATTTTACAGATTCTAACAGGCCTATTCCTAGCAATACACT    165
                     ||||||||||||||||||||||||||||||||||||||||||||||||||
C8_cytb-F2_20    151 CATTTGCTTAATTTTACAGATTCTAACAGGCCTATTCCTAGCAATACACT    200

A8_cytb-F1_20    166 ATACACCTGACACAACAACAGCATTCTCCTCTGTAACCCACATTTGCCGA    215
                     ||||||||||||||||||||||||||||||||||||||||||||||||||
C8_cytb-F2_20    201 ATACACCTGACACAACAACAGCATTCTCCTCTGTAACCCACATTTGCCGA    250

A8_cytb-F1_20    216 GACGTAAACTATGGCTGAATTATCCGATATATACACGCAAACGGGGCATC    265
                     ||||||||||||||||||||||||||||||||||||||||||||||||||
C8_cytb-F2_20    251 GACGTAAACTATGGCTGAATTATCCGATATATACACGCAAACGGGGCATC    300

A8_cytb-F1_20    266 AATATTTTTTATCTGCCTATTTATGCATGTAGGACGAGGCCTATACTATG    315
                     ||||||||||||||||||||||||||||||||||||||||||||||||||
C8_cytb-F2_20    301 AATATTTTTTATCTGCCTATTTATGCATGTAGGACGAGGCCTATACTATG    350

A8_cytb-F1_20    316 GATCATATACCTTCCTAGAAACATGAAACATCGGAGTAATCCTCCTATTT    365
                     ||||||||||||||||||||||||||||||||||||||||||||||||||
C8_cytb-F2_20    351 GATCATATACCTTCCTAGAAACATGAAACATCGGAGTAATCCTCCTATTT    400

A8_cytb-F1_20    366 GCGACAATAGCCACAGCATTCATAGGCTATGTTTTACCATGAGGACAAAT    415
                     ||||||||||||||||||||||||||||||||||||||||||||||||||
C8_cytb-F2_20    401 GCGACAATAGCCACAGCATTCATAGGCTATGTTTTACCATGAGGACAAAT    450

A8_cytb-F1_20    416 ATCATTCTGAGGAGCAACAGTTATTACCAACCTCCTTTCAGCAATTCCAT    465
                     ||||||||||||||||||||||||||||||||||||||||||||||||||
C8_cytb-F2_20    451 ATCATTCTGAGGAGCAACAGTTATTACCAACCTCCTTTCAGCAATTCCAT    500

A8_cytb-F1_20    466 ATATTGGCACAAACCTAGTCGAATGAATCTGGGGAGGATTCTCAGTAGAC    515
                     ||||||||||||||||||||||||||||||||||||||||||||||||||
C8_cytb-F2_20    501 ATATTGGCACAAACCTAGTCGAATGAATCTGGGGAGGATTCTCAGTAGAC    550

A8_cytb-F1_20    516 AAAGCTACCCTCACCCGATTTTTCGCCTTTCACTTTATTTTCCCATTCAT    565
                     ||||||||||||||||||||||||||||||||||||||||||||||||||
C8_cytb-F2_20    551 AAAGCTACCCTCACCCGATTTTTCGCCTTTCACTTTATTTTCCCATTCAT    600

A8_cytb-F1_20    566 CATCGCAGCCCTCGCCATAGTTCACCTACTCTTCCTCCACGAAACAGGAT    615
                     ||||||||||||||||||||||||||||||||||||||||||||||||||
C8_cytb-F2_20    601 CATCGCAGCCCTCGCCATAGTTCACCTACTCTTCCTCCACGAAACAGGAT    650

A8_cytb-F1_20    616 CCAACAACCCCACAGGAATTCCATCGGACACAGATAAAATTCCCTTCCAC    665
                     ||||||||||||||||||||||||||||||||||||||||||||||||||
C8_cytb-F2_20    651 CCAACAACCCCACAGGAATTCCATCGGACACAGATAAAATTCCCTTCCAC    700

A8_cytb-F1_20    666 CCTTATTACACCATTAAAGACATCCTAGGTGCTATCCTACTAATCCTCAT    715
                     ||||||||||||||||||||||||||||||||||||||||||||||||||
C8_cytb-F2_20    701 CCTTATTACACCATTAAAGACATCCTAGGTGCTATCCTACTAATCCTCAT    750

A8_cytb-F1_20    716 CCTCATGCTACTAGTACTATTCACGCCTGACTTACTCGGAGACCCAGACA    765
                     ||||||||||||||||||||||||||||||||||||||||||||||||||
C8_cytb-F2_20    751 CCTCATGCTACTAGTACTATTCACGCCTGACTTACTCGGAGACCCAGACA    800

A8_cytb-F1_20    766 ACTACACCCCAGCAA--CCACTT-ACACTCCCCCTCACATC-AACCTGAA    811
                     |||||||||||||||  |||||| ||||||||||||||||| ||||||||
C8_cytb-F2_20    801 ACTACACCCCAGCAAACCCACTTAACACTCCCCCTCACATCAAACCTGAA    850

A8_cytb-F1_20    812 TGATACTTCCTATTTGCGTACGC    834
                     ||||||||
C8_cytb-F2_20    851 TGATACTT---------------    858
```

**S3 Fig**. Pairwise sequence alignment between the partial sequences of cytb gene of the ovine heart cell line passage 26 (A8_cytb-F1_20) and passage 59 (C8_cytb-F2_20).

```
########################################
# Program: needle
# Rundate: Sun 19 Jun 2022 21:04:33
# Commandline: needle
#    -auto
#    -stdout
#    -asequence emboss_needle-E20220619-210431-0683-65551073-p2m.aupfile
#    -bsequence emboss_needle-E20220619-210431-0683-65551073-p2m.bupfile
#    -datafile EDNAFULL
#    -gapopen 10.0
#    -gapextend 0.5
#    -endopen 10.0
#    -endextend 0.5
#    -aformat3 pair
#    -snucleotide1
#    -snucleotide2
# Align_format: pair
# Report_file: stdout
########################################

#=======================================
#
# Aligned_sequences: 2
# 1: E8_MTCR-F1_20220424_SAMPLE1-MGCOMP-28572437
# 2: G8_MTCR-F2_20220424_SAMPLEA-36136530
# Matrix: EDNAFULL
# Gap_penalty: 10.0
# Extend_penalty: 0.5
#
# Length: 858
# Identity:     215/858 (25.1%)
# Similarity:   220/858 (25.6%)
# Gaps:         583/858 (67.9%)
# Score: 343.0
#
#
#=======================================

E8_MTCR-F1_20      1 GWTCACCATGCCGCGTGAAACCAACAACCCGCTCAGCAGGGATCCCTCTT     50

G8_MTCR-F2_20      1 --------------------------------------------------      0

E8_MTCR-F1_20     51 CTCGCTCCGGGCCCATTAACTGTGGGGGTAACTATTTAATGAACTTTAAC    100

G8_MTCR-F2_20      1 --------------------------------------------------      0

E8_MTCR-F1_20    101 AGGCATCTGGTTCTTTCTTCAGGGCCATCTCATCTAAAATCGCCCACTCT    150

G8_MTCR-F2_20      1 --------------------------------------------------      0

E8_MTCR-F1_20    151 TTCCCCTTAAATAAGACATCTCGATGGA--------CTAATGACTAATCA    192
                                            |||||        ||        ||||
G8_MTCR-F2_20      1 -----------------------ATGGATCACGGGTCT--------ATCA     19

E8_MTCR-F1_20    193 GCCCATGCCTAACATAACTGTGG----TGTCATGCATTTGGTATTTTTTA    238
                     .||.||   ||||...:|..|||    |..|||||||||||:||.|||||
G8_MTCR-F2_20     20 CCCTAT---TAACCAGWCACTGGAGCTTTCCATGCATTTGGWATCTTTTA     66

E8_MTCR-F1_20    239 ATTTTTGGGGATGCTTGGACTCAGCTATGGCCGTCTGAGGCCCCGACCCG    288
                                         ||           ||||
G8_MTCR-F2_20     67 --------------------TC-----------TCTG-------------     72

E8_MTCR-F1_20    289 GAGCATGAATTGTAGCTGGACTTAACTGCATCTTGAGCATCCTCATAATG    338
                                ||  |||.||..|||..||||    :|
G8_MTCR-F2_20     73 -----------GT--CTGCACRCAACACCATC----RC------------     93

E8_MTCR-F1_20    339 GTAAGCATG--GACATAATATAATTAATGGTCACAGGACATATC-TGCTG    385
                        |:||||  |||                ||.||..||||..| |.|||
G8_MTCR-F2_20     94 ---ARCATGCTGAC----------------TCCCACCACATCCCGTCCTG    124

E8_MTCR-F1_20    386 TAT---CGTGCATTTATATATTCTTTTTCCCCCCTTCCCCTTAAATATTT    432
                     .||   |.|||.|||   .||||.|.:|||
G8_MTCR-F2_20    125 AATGCGCCTGCCTTT---GATTCCTAKTCC--------------------    151

E8_MTCR-F1_20    433 ATCACCATTTTTAACACGCTTCCCCCTAGATATTAATATAAATTTATCCC    482
                       .|||||                      |||||||            |
G8_MTCR-F2_20    152 --AACCAT----------------------TATTAAT------------C    165

E8_MTCR-F1_20    483 GC-CCTCAATAC--TCAAATTCATACTCCAACCGAAGTAAATATATAGGC    529
                     || ||    |||  ||||..||     |||                  ||
G8_MTCR-F2_20    166 GCACC----TACGTTCAATGTC-----CCA------------------GC    188

E8_MTCR-F1_20    530 ACCTGGGTCACATACATAACGCATAGTTAATGTAGCTTAAACTTAAAGCA    579
                     .||          .||||||
G8_MTCR-F2_20    189 CCC----------GCATAAC------------------------------    198

E8_MTCR-F1_20    580 AGGCACTGAAAATGCCTAGATGAGTCTACTGACTCCATGAACATATAGGT    629
                        ||||.|.||||..|         ||.|.|.||||||
G8_MTCR-F2_20    199 ---CACTAATAATGTGT---------TATTTAATCCATG-----------    225

E8_MTCR-F1_20    630 TTGGTCCCAGCCTTCCTGTTAACTTTCAATAGACTTATACATGCAAGCAT    679
                                   |.|||          .||||  ||||| .|||.||.
G8_MTCR-F2_20    226 --------------CTTGT----------AAGAC--ATACA-ACAACCAA    248

E8_MTCR-F1_20    680 CCACGCCCCGGTGAGTAACGCCCTTCGAATCACACAGGACTAAAAGGAGC    729
                     ||                              ||||..||||.||
G8_MTCR-F2_20    249 CC------------------------------CACAACACTACAA-----    263

E8_MTCR-F1_20    730 AGGTATCAAGCACACACTCTTGTAGCTCACAACGCCTTGCTTAACCACAC    779
                                |.|.|.||        .|.|||      ||||||||.||
G8_MTCR-F2_20    264 -----------AAAAATTC--------AAAAAC------CTTAACCAAAC    288

E8_MTCR-F1_20    780 CCCC---------ACGGGAGACAGCAGTAACAAAAATTAAGCCATAAACG    820
                     ||||         ||..
G8_MTCR-F2_20    289 CCCCCCCCCYTKWACCT---------------------------------    305

E8_MTCR-F1_20    821 AAAGTTTG    828

G8_MTCR-F2_20    306 --------    305
```

**S4 Fig.** Pairwise sequence alignment between the partial sequences of the ovine mitochondrial segments spanning part of the control region, the tRNA-Phe gene, and part og the 12S rRNA gene of heart cell line passage 26(E8_MTCR-F1_20) and passage 59 (G8_MTCR-F2_20).

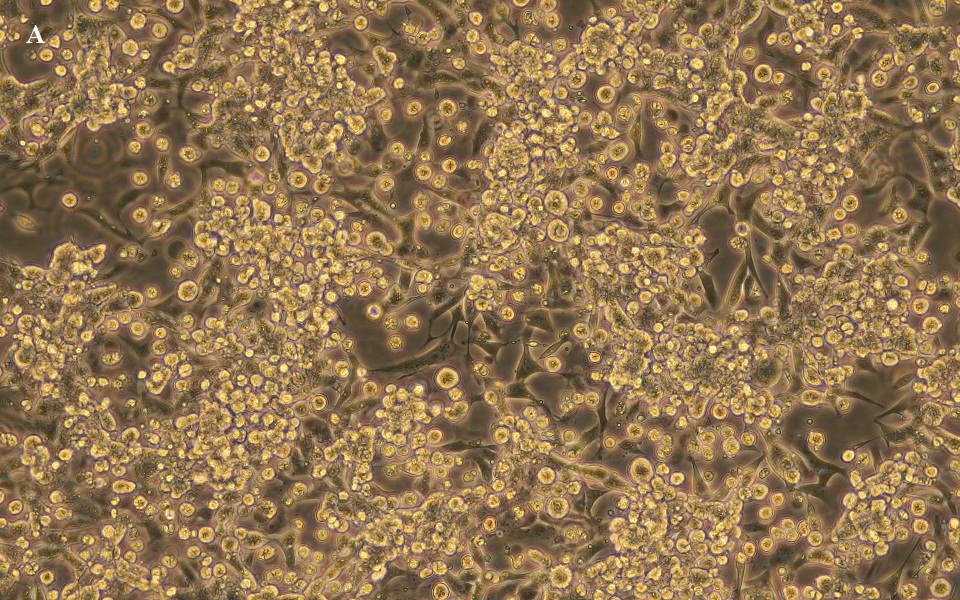

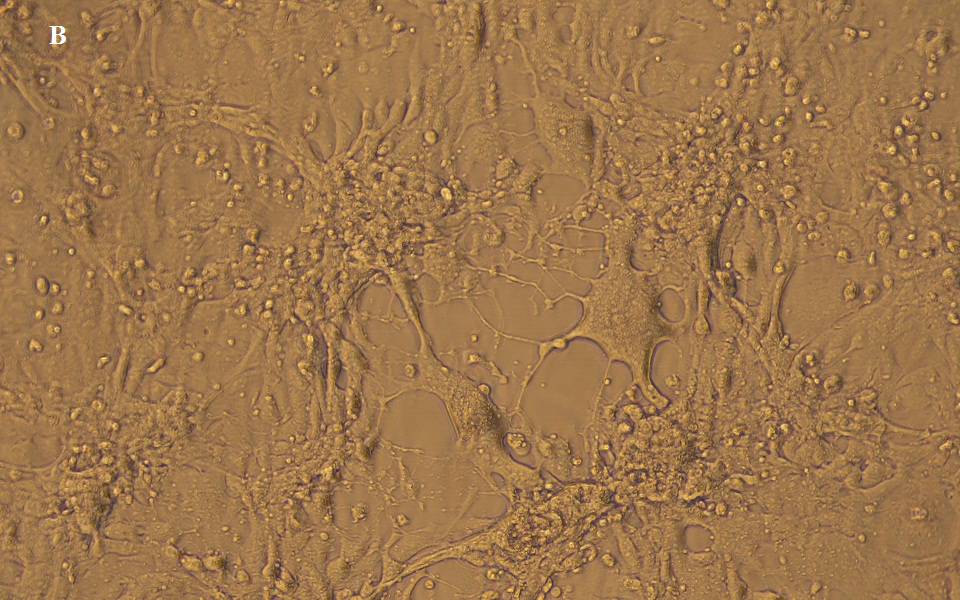

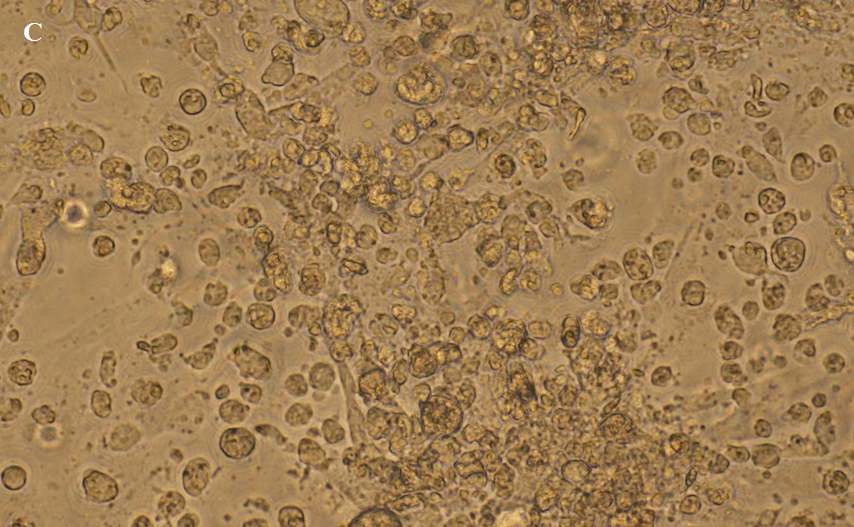

**S5 Fig**.Sensitivity of the FOH-SA cell line to viruses.a) Cytopathic effect (CPE) of sheeppox virus, b) CPE of camelpox virus, c) CPE of PPR virus, phase contrast X200.

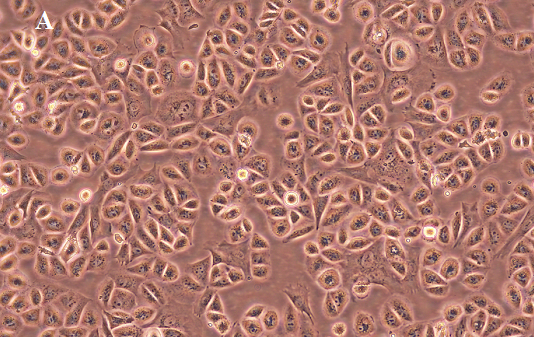

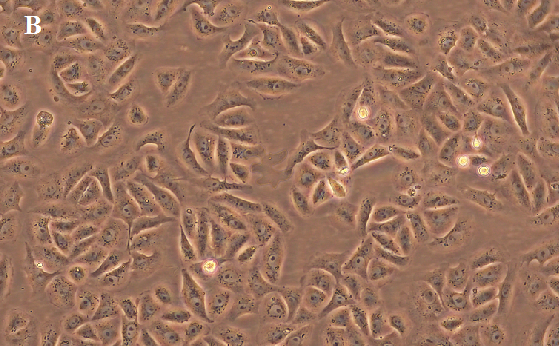

**S6 Fig**. The morphology of a three hour culture of a) FOH-SA line passage 55 and b) Vero ATCC CCL-81cell line, phase contrast X200.

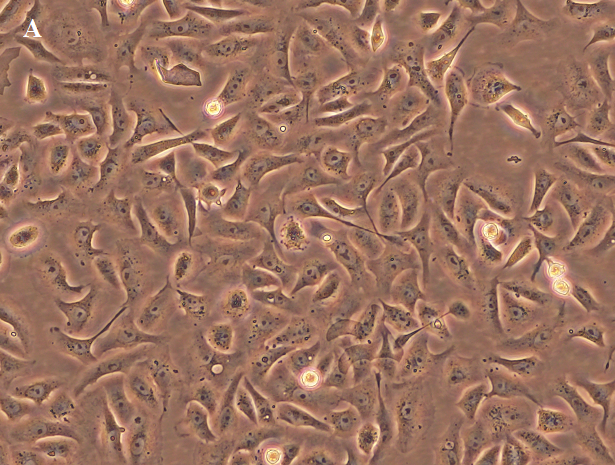

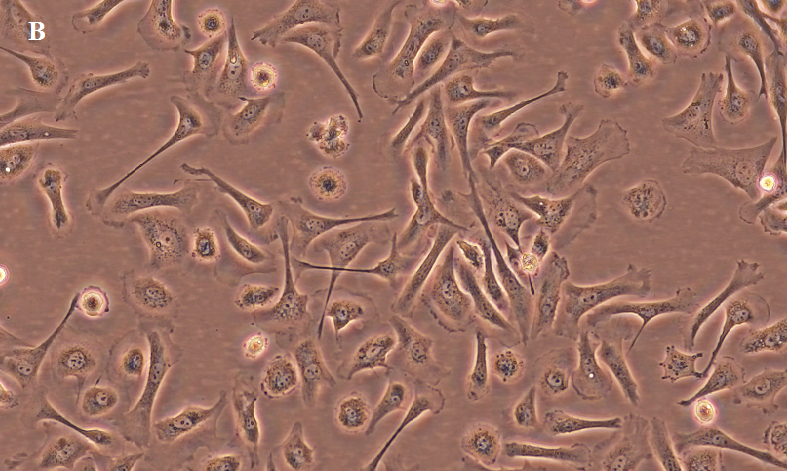

S7 Fig. a) Morphology of a three hour culture of Vero ATCC CCL-81 cells and b) morphology of a three hour culture of the ovine heart cell line passage 22, Both cell lines show epithelial-like cells and fibroblast-like cells with threads, phase contrast X200.

S1Table. Successive SNPs that were called inDNA of cells before cell fusion and were not called in DNA of cells after cell fusion and the genes spanning the areas between the SNPs.

| **Genes** | **Location** | **Chromosome** | **SNP name** | **SNP index** |
| --- | --- | --- | --- | --- |
| *ADAMTS19* | 23286278  23332486  23340233  23352342  23400692 | 5 | OAR5_23286278.1  OAR5_23332486.1  OAR5_23340233.1  OAR5_23352342_X.1  OAR5_23400692.1 | 28172  28173  28174  28175  28176 |
| *ESRRB* | 91618922  91628108  91692431  91765475  91823711 | 7 | OAR7_91618922_X.1  OAR7_91628108.1  OAR7_91692431.1  OAR7_91765475.1  OAR7_91823711.1 | 32826  32827  32828  32829  32830 |
| *ACSS2, MYH7B, TRPC4AP, EDEM2,*  *PROCR, MMP24,*  *EIF6, FAM83C*, *UQCC1* | 67008189  67067820  67246527  67400889  67425309 | 13 | OAR13_67008189.1  OAR13_67067820.1  OAR13_67246527.1  OAR13_67400889.1  OAR13_67425309.1 | 8579  8580  8581  8582  8583 |
| *PRKAA2, CBA* | 32984769  33007740  33043220  33161254 | 1 | OAR1_32984769.1  OAR1_33007740.1  OAR1_33043220.1  OAR1_33161254.1 | 3697  3698  3699  3700 |
| *SMARCA1* | 120910363  120998827  121015458  121076593 | X | OARX_120910363.1  OARX_120998827.1  OARX_121015458.1  OARX_121076593.1 | 36429  36430  36431  36432 |
| *GALNT13* | 167547154  167611214  167667143 | 2 | OAR2_167547154.1  OAR2_167611214.1  OAR2_167667143.1 | 15421  15422  15423 |
| *BICD1* | 195730138  195768950  195793605 | 3 | OAR3_195730138.1  OAR3_195768950.1  OAR3_195793605_X | 23918  23919  23920 |

S1 Table .*Continued*

| **Genes** | **Location** | **Chromosome** | **SNP name** | **SNP index** |
| --- | --- | --- | --- | --- |
| *PTPRO* | 214578400  214663567  214698761 | 3 | OAR3_214578400.1  OAR3_214663567.1  OAR3_214698761.1 | 24267  24268  24269 |
| *CAMKMT* | 84974488  85000557  85042665 | 3 | OAR3_84974488.1  OAR3_85000557.1  OAR3_85042665.1 | 25529  25530  25531 |
| *ARHGAP26* | 56102371  56257273  56291790 | 5 | OAR5_56102371.1  OAR5_56257273.1  OAR5_56291790.1 | 28646  28647  28648 |
| *GRIA1* | 67605574  6761259  67649257 | 5 | OAR5_67605574.1  OAR5_6761259.1  OAR5_67649257.1 | 28784  28785  28786 |
| *CLVS1, ASPH* | 43100350  43198107  4322683 | 9 | OAR9_43100350.1  OAR9_43198107.1  OAR9_4322683.1 | 35082  35083  35084 |
| *AMER2, C1q and TNF related 9, MIPEP* | 35950338  36265107  36302356 | 10 | OAR10_35950338.1  OAR10_36265107.1  OAR10_36302356.1 | 5236  5237  5238 |
| *BTRC* | 24747565  24756524  24815596 | 22 | OAR22_24747565.1  OAR22_24756524.1  OAR22_24815596.1 | 19557  19558  19559 |
| *ABRAXAS2,*  *ZRANB1, CTBP2* | 47793640  48066306  4815146 | 22 | OAR22_47793640.1  OAR22_48066306.1  OAR22_4815146.1 | 19893  19894  19895 |
| *KIFBP, SUPV3L1,*  *VPS26A, HKDC1*  *SRGN* | 26224691  26553035  26619489 | 25 | OAR25_26224691.1  OAR25_26553035.1  OAR25_26619489.1 | 21426  21427  21428 |

ADAM metallopeptidase with thrombospondin type 1 motif 19(*ADAMTS19*), estrogen related receptor beta(*ESRRB*), Acyl-CoA synthetae short chain family member 2(*ACSS2*), myosin heavy chain 7B(*MYH7B*), transient receptor potential cation channel subfamily C member 4 associated protein(*TRPC4AP*), -ER degradation enhancing alpha-mannosidase-like protein2(*EDEM2*), protein C receptor(*PROCR*), matrix metallopepridase 24(*MMP24*), eukaryotic translation intiation factor 6(*EIF6*), family with sequence similarity 83 member C(*FAM83C*), ubiquinol-cytochrome C reductase complex assembly factor 1(*UQCC1*), protein kinase AMP-activated catalytic subunit alpha 2 (*PRKAA2*), complement C8 alpha chain(*CBA*), SW1/SNF related, matrix associated, actin dependent regulator of chromatin, subfamily a, member 1(*SMARCA1*), polypeptide N-acetylgalactosaminyltransferase 13(*GALNT13*), BICD cargo adaptor 1(*BICD1*), protein tyrosine phosphatase receptor type O(*PTPRO*), calmodulin-lysin N-methyltransferase (*CAMKMT*) Rho GTpase activating protein 26(*ARHGAP26*), glutamate ionotropic receptor AMPA type subunit 1(*GRIA1*), aspartate beta-hydroxylase(*ASPH*), clavesin(*CLVS1*), APC membrane recruitment protein(*AMER2*), mitochondrial intermembrane peptidase(*MIPEP*), abraxas 2, BRISC complex subunit (*ABRAXAS2*), zinc finger RANBP2-type containing 1(*ZRANB1*), C-treminal binding protein 2(*CTBP2*)beta-transducin repeat containing E3 ubiquitin protein ligase(*BTRC*), kinesin family binding protein(*KIFBP*), Suv3 like RNA helicase(*SUPV3L1*),VPS26, retromer complex component A(*VPS26A*), hexokinase domain containing 1(*HKDC1*), serglycin(*SRGN*).
